# Aberrant replication licensing drives Copy Number Gains across species

**DOI:** 10.1101/2020.10.10.334516

**Authors:** Patroula Nathanailidou, Michalis Petropoulos, Styliani Maxouri, Eirini Kasselimi, Ioanna Eleni Symeonidou, Ourania Preza, Iris Spiliopoulou-Sdougkou, Vladimir Beneš, Stavros Taraviras, Zoi Lygerou

## Abstract

Copy Number Gains (CNGs) lead to genetic heterogeneity, driving evolution and carcinogenesis. The mechanisms promoting CNG formation however remain poorly characterized. We show that abnormal expression of the replication licensing factor Cdc18 in fission yeast, which leads to genome-wide re-replication, drives the formation of CNGs at different genomic loci, promoting the acquisition of new selectable traits. Whole genome sequencing reveals Mb long, primarily extrachromosomal amplicons. Genetic analysis shows that homology-mediated repair is required to resolve re-replication intermediates into heritable CNGs. Consistently, we show that in mammalian cells overexpression of CDC6 and/or CDT1 leads to CNGs and promotes drug resistance. In human cells, multiple repair pathways are activated upon rereplication and act antagonistically, with RAD52 promoting and 53BP1 inhibiting CNG formation. In tumours, CDT1 and/or CDC6 overexpression correlates with copy number gains genome-wide. We propose re-replication as an evolutionary-conserved driver of CNGs, highlighting a link between aberrant licensing, CNGs and cancer.

---

Copy Number Gains (CNGs) drive genome evolution, leading to phenotypic variation and promoting species and cancer evolution^1,2^. In tumorigenesis, extra copies of oncogenes advance malignant transformation and provide a proliferative advantage for clonal expansion^2–4^, while amplification of genes related to drug metabolism result in acquired resistance to cancer treatment^5^. Despite the importance of CNGs for multiple biological processes, mechanisms promoting their generation remain poorly understood. Non-allelic homologous recombination and microhomology mediated pathways including Break-induced replication and single-strand annealing contribute to CNGs in meiotic and mitotic cells^6^ while Break-Fusion-Bridge (BFB) cycles promote gene amplification in cancer cells^7,8^. Re-Replication has also been proposed as a putative mechanism contributing to CNGs^9,10^. Cells normally ensure that each part of their genome is replicated once and only once per cell cycle through multiple control mechanisms^11^. Aberrations in these controls in different species leads to rereplication^12–16^,the generation of multiple copies of DNA, arranged as replication bubbles within bubbles. The fate of these aberrant replication intermediates however remains elusive. In *Saccharomyces cerevisiae*, where limited rereplication is observed^13^, re-firing of a specific origin of replication can generate intra-chromosomal, tandemly arrayed copies of this locus^14,17^. Can rereplication lead to copy number gains at multiple genomic loci and in different species and could it therefore constitute an evolutionarily conserved driver of genome plasticity? By combining experiments in fission yeast, where rereplication can be induced at multiple locations across the genome^18,19^ and mammalian cells, we show that re-replication promotes the generation of CNGs in different genomic loci and can enhance the acquisition of new selectable traits. We show that the resolution of re-replication intermediates into stable amplicons is channelled through homology and microhomology-mediated repair mechanisms. Whole genome sequencing in fission yeast reveals Mb long, extrachromosomal amplicons. Deregulated replication licensing is present from the initial steps of cancer development^9,20^. We show that cancer samples with deregulated licensing show increased CNGs genome-wide, indicating clinical relevance for re-replication induced CNGs. Our work highlights re-replication induced by aberrant licensing control as an evolutionarily conserved mechanism for CNGs.

## Results

### Re-replication promotes CNGs at the *sod2* locus in fission yeast

To test whether re-replication induction drives CNGs we employed fission yeast strains carrying a *cdc18* variant (*d55P6*, see Online Methods), under the inducible promoter *nmt1*. In fission yeast, overexpression of the licensing factor Cdc18 is sufficient to bring about re-replication across the genome^18,19^. We initially analysed the genomic locus *sod2*, known to confer resistance to lithium chloride (LiCl) when present in multiple copies. Following rereplication induction, cells were plated under LiCl selection. The incidence of *sod2* amplification was assessed by the number of colonies arising under selection corrected to viability (Extended Data Fig. 1a) and compared to the control non-re-replicating condition. Re-replication greatly enhances the incidence of *sod2* amplification (Fig. 1a). qRT-PCR analysis confirmed that LiCl resistance results from multiple copies of the *sod2* gene in rereplicating (Fig. 1b) and control (Extended Data Fig. 1b) conditions. The result was verified in multiple induction time-points (Extended Data Fig. 1c). We conclude that re-replication promotes CNG formation at the *sod2* locus.

**Fig. 1.**
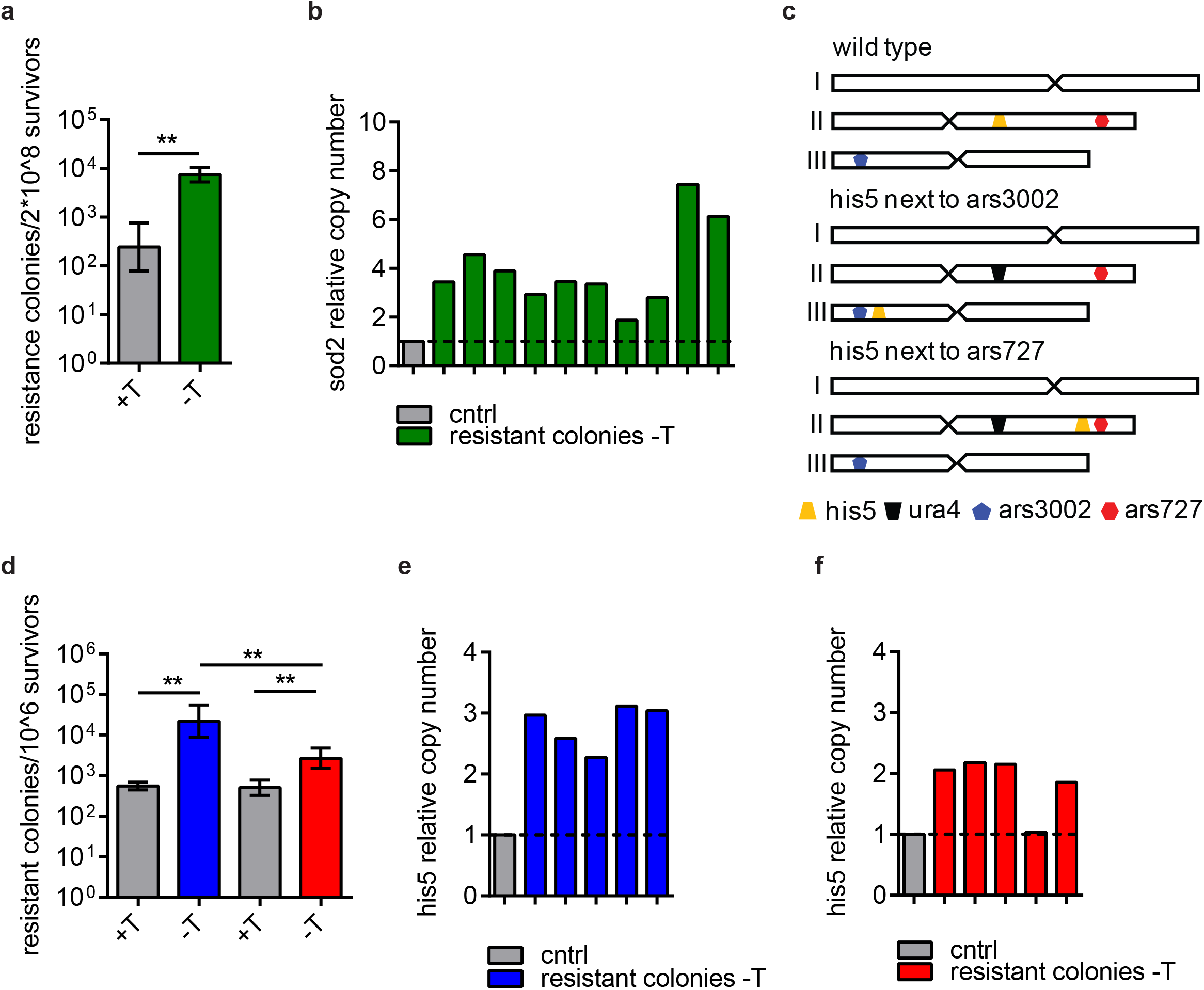
Re-replication promotes CNG formation at different genomic positions in fission yeast, in an origin activity dependent manner. **a**, *nmt1-d55P6-cdc18* cells were cultured in the absence or presence of thiamine for 23 hours, to induce re-replication or not, respectively. In each condition, 2×10^8^ cells were plated in the presence of 40mM LiCl and thiamine and the number of LiCl resistant colonies, corrected to survival, is presented; *n=5* independent biological replicates. **b**, Resistant colonies were cultured in the presence of thiamine in non-selective medium and genomic DNA was isolated. The relative copy number of the *sod2* locus upon re-replication was determined by qRT PCR. **c**, Schematic representation of the genotypes of the strains constructed. The endogenous *his5* gene was deleted using a *ura4* cassette. *his5* with its regulatory elements was integrated next to an efficient (ars3002) or dormant (ars727) origin. **d**, Re-replication was induced, or not, for 25 hours for both strains constructed and 10^6^ cells were plated under non-re-replicating conditions in the presence of 5mM 3-AT. The corrected numbers of resistant colonies arising upon re-replication, or not, are presented; *n*=*6* independent biological replicates. **e**, **f**, The relative copy number of the *his5* locus was determined upon re-replication in strains carrying the *his5* locus next to ars3002 (e) and ars727 (f), using qRT PCR. The non-parametric, two-tailed Mann-Whitney U test was used to assess the significance of differences in each case.

### Re-replication induced CNGs occur at multiple loci and are affected by local origin activity

To show that our results are not locus or drug specific and to uncover possible position effects, an alternative CNG assay was set up, based on the *his5* locus^21^ and its ability to confer resistance to 3 amino-triazole (3-AT), when present in multiple copies. In a *nmt1-d55P6*-*cdc18* background strain, the endogenous *his5* locus was deleted and *his5* was ectopically integrated close to an efficient (ars3002) or a dormant (ars727) origin (Fig. 1c). 3AT resistance-based assays were performed for each integration site following re-replication induction. 5 mM 3AT were sufficient to select multiple versus single *his5* copies (Extended Data Fig. 1d, e, f, g). Increased incidence of *his5* CNGs upon re-replication was observed in both cases (Fig. 1d). However, more resistant colonies were formed when the *his5* cassette was integrated next to the efficient origin compared to the dormant (Fig. 1d). qPCR verified that increased 3AT resistance rates were accompanied by extra *his5* copies in both integration sites (Fig. 1e, f, Extended Data Fig. 1h, i), with higher copies observed at the vicinity of the efficient origin. We conclude that re-replication promotes formation of CNGs at different genomic positions, and under different selection and that the frequency of these events is positively correlated to the activity of nearby origins.

### Re-replication generates Mb-long mostly extrachromosomal CNGs

To elucidate the genomic structures generated through re-replication, Whole Genome Sequencing (WGS) was performed in three 3-AT and four LiCl resistant colonies and Copy Number was analyzed across the genome (Online Methods) (Fig. 2a, c, d and Extended Data Fig. 2a, b).

**Fig. 2.**
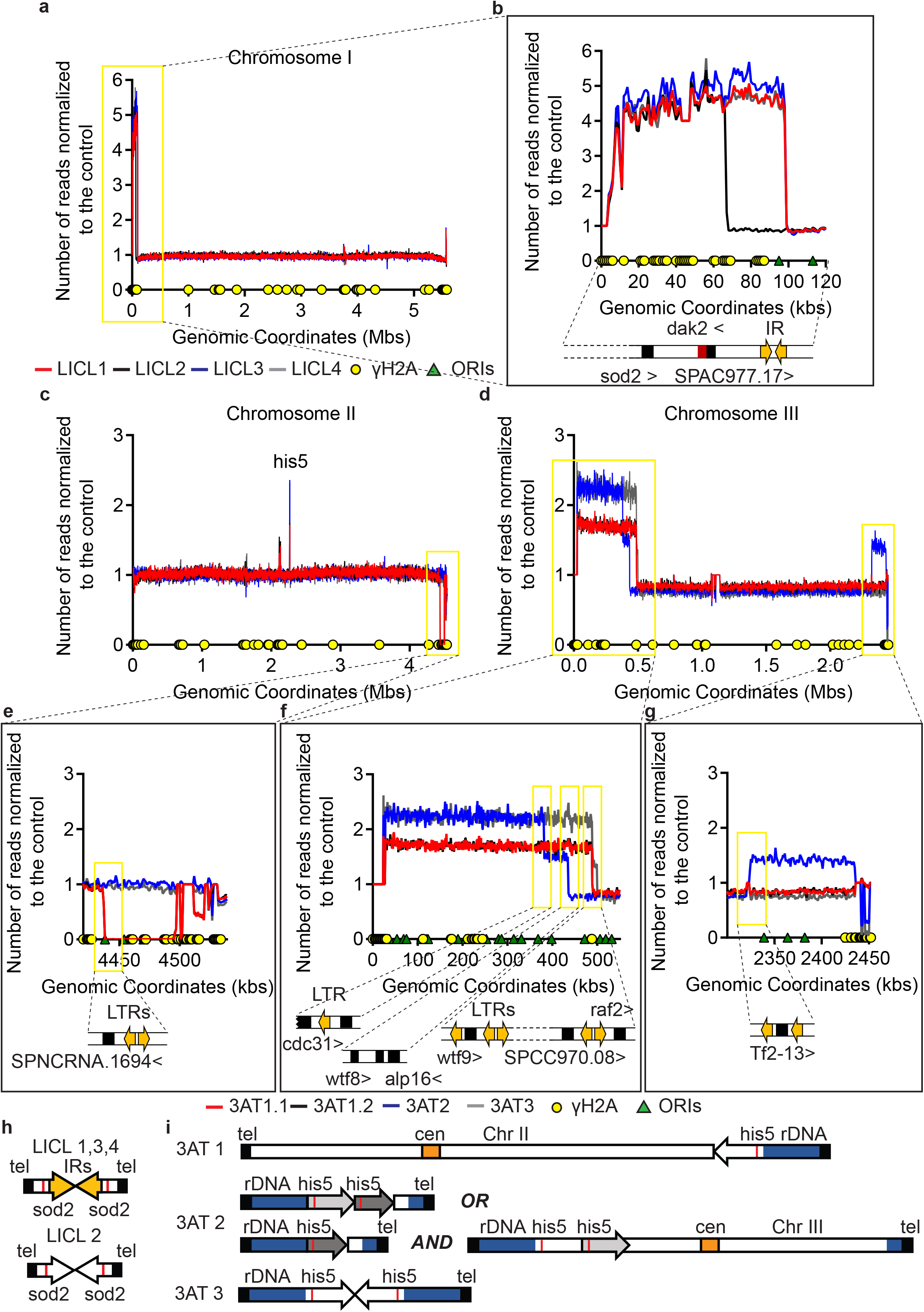
Re-replication induces mostly extrachromosomal, Mb-long CNGs in fission yeast. LiCl and 3-AT resistant colonies were cultured under 20 mM LiCl and 5 mM 3-AT selection, respectively, in non-re-replicating conditions and genomic DNA was isolated. Samples from two non-resistant control strains were also prepared. Whole Genome Sequencing was performed in all samples. Total reads per 1000 base pairs in each resistant colony were normalized to the total reads per 1000 base pairs of the non-resistant control and plotted. γH_2_A enriched sites, indicating Common Fragile Sites (CFS), are depicted as yellow circles, and replication origins are shown as green triangles. **a**, **c**, **d**, The observed alterations are highlighted in yellow boxes for LiCl and 3AT resistant colonies, respectively. **b**, **e**, **f**, **g**, Detailed view of the amplified regions in LiCl and 3AT samples. The genomic features underlying the borders of the amplicon are presented. Amplicon borders coincide with Long Terminal Repeats (LTRs) and transposable elements. **h**, **i**, The exact structures of amplicons were revealed after sequencing data analysis and are presented for LiCl and 3AT, respectively.

In LiCl resistant colonies, the genome was overall intact except for the telomere proximal region on the left arm of Chr. I, where the sod2 locus resides, which was amplified in all samples tested (Fig. 2a, Extended Data Fig. 2a). Sequencing revealed a recurrent amplicon (LiCl 1, 3, 4), with an inverted repeat located telomere-proximal to the break point of the amplicon (Fig. 2b). A shorter amplicon (LiCl 2), ending in a Common Fragile Site (CFS) indicated by γH2A enrichment^22^, was also observed (Fig. 2b). Structural analysis showed that CNGs are present as extrachromosomal inverted repeats (Fig. 2h). Observed genotypes were verified by Pulsed Field Gel Electrophoresis (PFGE) in more clones. Amplicons appeared as extra bands of 180 kb or 225 kb, confirming the structures predicted by WGS and indicating extrachromosomal amplicons with telomeres (Extended Data Fig. 2c, d), consistent with earlier work on spontaneous *sod2* amplification^23^. 30% of samples present extra copies as stable integrates in Chr. I, indicated by the shifted band of Chr. I (Extended Data Fig. 2c). Abnormal size of Chr. III indicates copy number alterations in the rDNA repeats upon re-replication^24^.

For the 3-AT isolates, the main genomic region affected resides at the left arm of Chr. III, where a 0.5 Mb amplicon is observed encompassing the ectopically integrated his5 selection cassette (Fig. 2d). Structural analysis is consistent with tandem and inverted extrachromosomal repeats (Fig. 2i). In one of the clones (analysed in duplicate, 3AT1.1 and 3AT1.2) the amplified region on the left arm of Chr. III is accompanied by a terminal deletion at the right arm of Chr. II, consistent with a chromosomal translocation of the additional copy (Fig. 2i). Amplicon boundaries coincide with Long Terminal Repeats (LTRs) and transposable elements (Fig. 2e, f, g). The homology shared between these repetitive elements (>200 bp), points to homology-mediated resolution of the structures emerging upon re-replication at this region. PFGE revealed frequent loss of Chr. III (Extended Data Fig. 2c) and 3AT amplicon visualization was not possible, consistent with rDNA repeat imbalances^24^, which produce a range of amplicon sizes. PFGE followed by Southern blotting against the *his5* gene verified the expected smeary pattern (Extended Data Fig. 2e), while the presence of large amplicons (2-5,7 Mb) supports structures containing rDNA repeats and telomeres (Extended Data Fig. 2e).

Taken together, WGS reveals mostly extrachromosomal long amplicons while the location of the breakpoint centromere proximal to the selection gene is consistent with a BFB-independent amplification mechanism.

### Formation of re-replication induced CNGs is channeled through homology-mediated pathways and requires an intact checkpoint mechanism in fission yeast

Next, the downstream molecular mechanisms required for formation of CNGs upon re-replication were explored. A set of strains bearing a deletion of core DNA damage repair components, each inactivating a specific repair pathway, was constructed in a *d55P6-cdc18* overexpressing background. Comparable re-replication levels between wild type and mutant strains were induced (Extended Data Fig. 3a-e) and colony resistance assays were performed both for the *sod2* and the *his5* locus. *pku70* and *rad22A* mutants were used as indicators of Non-Homologous End Joining (NHEJ) and Homologous Recombination (HR), respectively^25^. The number of resistant colonies remained unaffected upon *pku70* deletion for both loci tested (Fig. 3a, 3f), suggesting that the resolution of re-replication intermediates to CNGs is not channeled through NHEJ. While the effect of *rad22A* deletion at the *his5* locus could not be assessed (see Online Methods), *rad22A* deletion caused a major reduction of LiCl resistant colonies (Fig. 3b), pointing to HR-related mechanisms. To further examine the HR sub-pathways mediating CNG formation at both loci, specific mutants inactivating the classical HR (Rhp51), the Break Induced Replication (BIR, *cdc27-D1*) and the alternative Microhomology Mediated End Joining (MMEJ) pathway (*rad16Δ*)^26,27^ were tested. CNGs at the *sod2* locus proceeds in a *rhp51* independent manner (Fig. 3c), pointing to the alternative MMEJ pathway for resolution. Consistently, the MMEJ specific mutant *rad16Δ* presents decreased frequency of CNG formation at the *sod2* locus (Fig. 3e), while the BIR specific mutant *cdc27-D1*^28^ has no effect (Fig. 3d). On the other hand, *his5* CNGs present Rhp51 dependency (Fig. 3g), suggesting a classical HR or BIR pathway for resolution. BIR inactivation increases the incidence of *his5* CNGs (Fig. 3h), indicating an antagonistic action of this pathway and pointing to classical HR for the formation of the *his5* CNGs. The *rad16* mutant does not present any effect (Fig. 3i), as expected. Collectively, unstable intermediates caused upon re-replication induction are stabilized through homology mediated mechanisms, while locus characteristics determine the genetic requirements for the resolution and affect the specific pathway utilized at each locus.

**Fig. 3.**
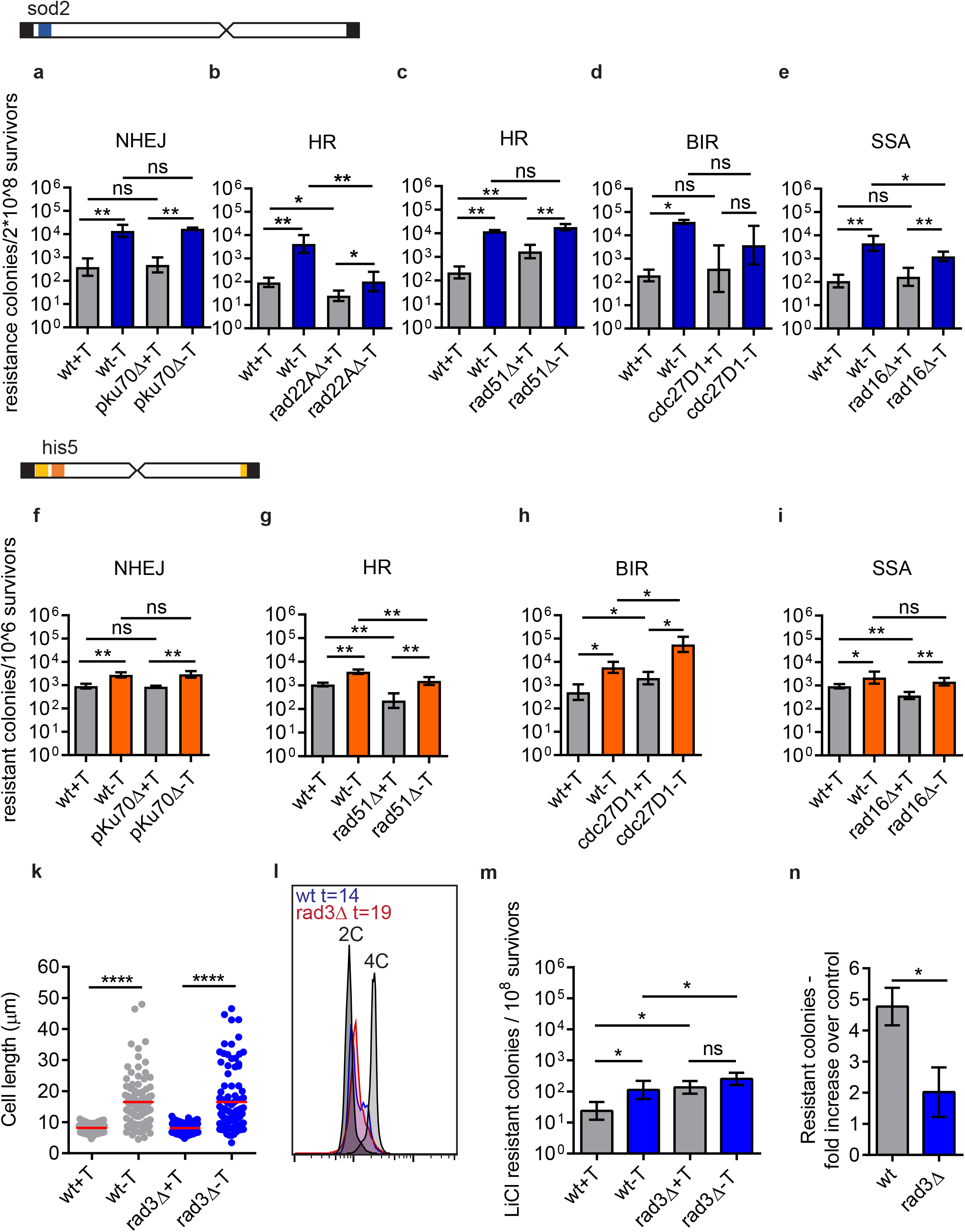
Re-replication induced CNGs are stabilized through homology and microhomology-mediated repair pathways and their formation is affected by checkpoint activation. *nmt1-d55P6-cdc18* strains deleted or mutated for DNA damage repair factors, acting in core repair pathways, were constructed. **a**, **f**, *pKu70Δ* **b**, *rad22AΔ* (Rad52) and **c**, **g**, *rhp51Δ* (Rad51) **d**, **h**, *cdc27-D1* (Polδ) and **e**, **i**, *rad16Δ* mutants were used to inactivate NHEJ, HR, BIR and MMEJ, respectively. Wild type and mutant cells were cultured in parallel in the presence or absence of thiamine. The time interval of the induction was adjusted to achieve same re-replication levels between wild type and mutant strains each time. 2×10^8^ cells were counted and plated in the presence or absence of 40 mM LiCl or 5 mM 3AT under non-re-replicating conditions. The number of resistant colonies upon re-replication, or not, was counted both for wild type and mutant strains. The numbers were corrected to survival and plotted; *n*=*5* independent biological replicates were performed for *pKu70Δ*, *rad22AΔ* (Rad52), *rhp51Δ* (Rad51), *rad16Δ* strains and *n*=*4* biological replicates for the *cdc27-D1* (Polδ) mutant**. k**, Cell length analysis in cell strains overexpressing the wild type Cdc18 (*nmt1-cdc18*), in the presence (wt) or absence (*rad3Δ*) of Rad3. Re-replication was induced for 21 hours, or not, and cells were fixed with PFA. Microscopy pictures were taken and the cell length of 100 cells per specimen was measured using ImageJ. Wild type Cdc18 is able to arrest the cell cycle upon re-replication induction, indicated by the typical elongated phenotype. **l**, FACS analysis for the *nmt1-cdc18* strain. Overexpression of Cdc18 was induced for 14 hours in the wt and 19 hours in the *rad3Δ* cells. The respective re-replication levels were monitored using PI staining and FACS analysis and the results are presented. These timepoints were used for the following colony formation assays. **m**, Re-replication was induced or not, in wild type and *rad3Δ nmt1-cdc18* cells. 2×10^8^ cells were plated in the presence of 40mM LiCl under non-re-replicating conditions and the number of LiCl resistant colonies upon re-replication, or not, corrected to survival, is presented. **n**, The number of colonies in the re-replicating condition for both strains is also presented as fold change over the control, non-re-replicating condition. The non-parametric, two-tailed Mann-Whitney U test was used to assess the significance of differences in each case.

We then examined the effect of checkpoint activation on re-replication induced CNGs, exploiting a cell strain lacking the central sensor kinase Rad3 (*rad3Δ*). We were not able to detect overt re-replication upon overexpression of the N-terminally truncated *d55P6 cdc18* in a *rad3Δ* background (Extended Data Fig. 3f, g), as cells were unable to arrest their cell cycle in the absence of Rad3 and entered aberrantly into mitosis (Extended Data Fig. 3h), in line with earlier work^29^. Thus, wild type *cdc18*, which is able to arrest the cell cycle upon re-replication in the absence of Rad3 (Fig. 3k) by directly inhibiting the mitotic CDK^29^, was overexpressed in a wild type or *rad3Δ* background and LiCl resistant colonies were quantified following re-replication (Fig. 3l, m, n). While Rad3 deletion increases the number of spontaneous CNGs under normal conditions, as previously reported^30^ (Fig. 3m and Extended Data Fig. 3i), a reduced number of re-replication induced resistant colonies are observed when Rad3 is absent (Fig. 3m, n), suggesting that checkpoint activation is important for CNGs upon re-replication induction.

### DNA re-replication promotes CNGs in mammalian cells

To examine whether re-replication induced CNGs are observed in mammalian cells, we took advantage of the *DHFR* locus and the methotrexate (MTX) resistance acquired by multiple copies of the gene ^31^. Re-replication was induced, through CDT1 and CDC6 overexpression in U2OS and Saos-2 cancer cells and the frequency of re-replication induced *DHFR* CNGs was examined based on the number of the MTX resistant colonies formed. MTX toxicity in U2OS and Saos-2 cell lines was evaluated by MTT assay (Extended Data Fig. 4a). The number of MTX resistant colonies was corrected to survival following CDT1 and CDC6 overexpression (Extended Data Fig. 4b). More MTX resistant colonies arose in U2OS and Saos-2 cells overexpressing CDT1 and CDC6 compared to control transfected cells (Fig. 4a, b). qRT-PCR verified that MTX resistant clones from re-replicating or control cells exhibited a higher copy number compared to non-resistant cells (Fig. 4c and Extended Data Fig. 4c). Therefore, cancer cells with aberrant expression of CDT1 and CDC6 acquire CNGs at the *DHFR* genetic locus. To test whether this effect is also present in normal cells, similar experiments where performed in NIH3T3 cells stably overexpressing Cdt1 (NIH3T3 Cdt1) (Extended Data Fig. 4d, e)^32^. NIH3T3 Cdt1 expressing cells form more MTX resistant colonies compared to control NIH3T3 cells, confirming our results (Fig. 4d). Combined overexpression of Cdt1 and Cdc6 increased further the incidence of *Dhfr* CNGs, while Cdc6 overexpression alone had no effect (Fig. 4d). qRT-PCR in MTX resistant survivors revealed *Dhfr* CNGs both in NIH3T3 Cdt1 and NIH3T3 MSCV cell lines (Fig. 4e, Extended Data Fig. 4f). Notably, Cdt1 overexpression lead to DNA double strand breaks, marked by phosphorylation of H2AX (Extended Data Fig. 4h, i) and chromosomal instability (data not shown). Overall, these results suggest that DNA re-replication induces CNGs in different cancer and normal mammalian cells.

**Fig. 4.**
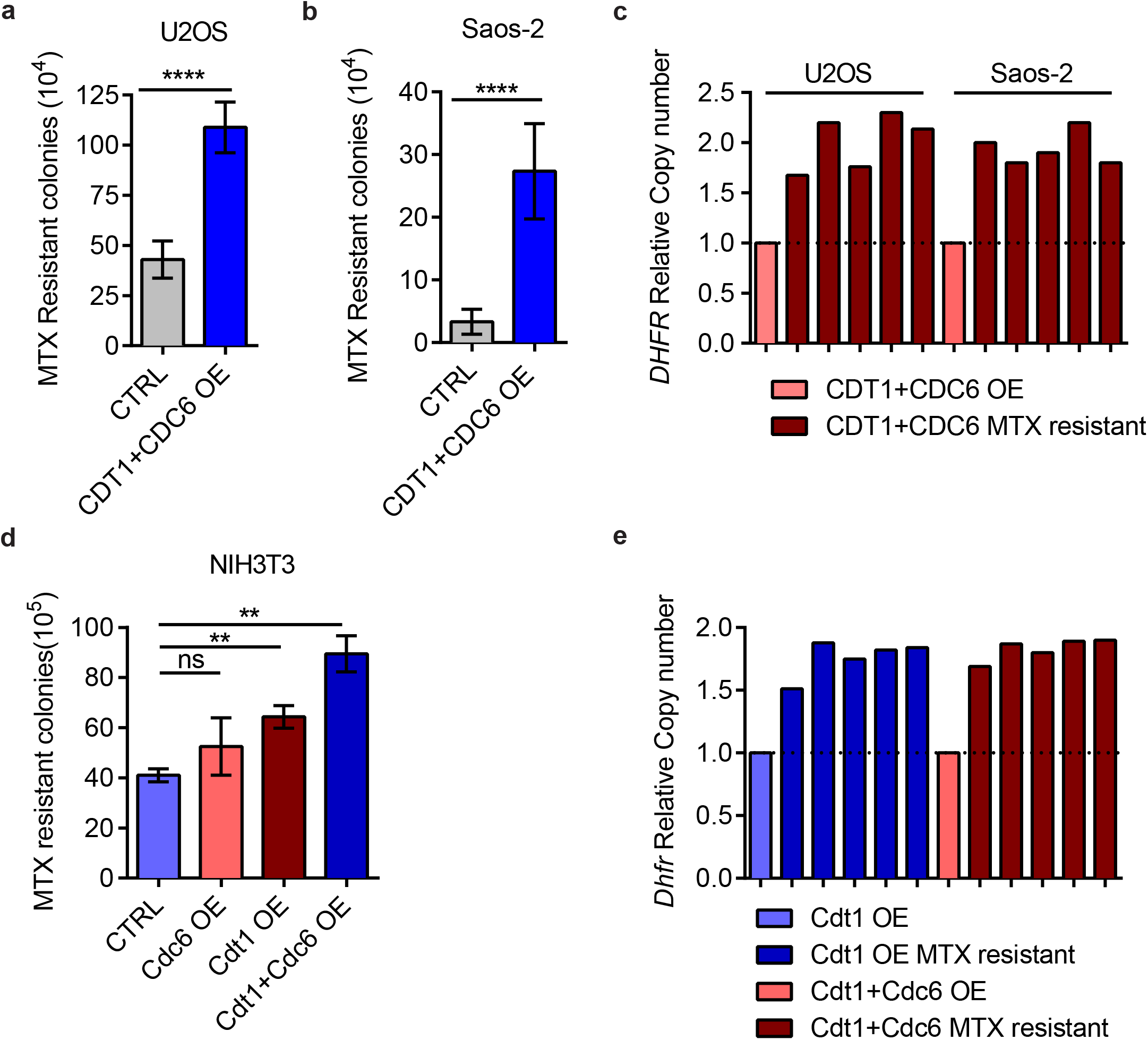
Ectopic expression of CDT1 and CDC6 enhances the frequency of *DHFR* CNGs in mammalian cells. **a,** Number of MTX resistant colonies upon re-replication, or not, in U2OS and **b,** Saos-2 cells, corrected to survival. Human U2OS and Saos-2 parental cells were transiently transfected with CDT1-GFP and Cherry-CDC6 expression vectors or GFP and Cherry as control, respectively, and after 48 hours, 10^4^ cells were seeded in 10 cm cell culture plates. After 24 hours cells were incubated in the presence of MTX for 20 days. After 20 days, plates were fixed, and resistant colonies were stained with methylene blue and counted; *n*=3 biologically independent replicates, CTRL vs CDT1+CDC6, ****P< 0.0001, statistical analysis was performed using two-tailed Mann-Whitney *U-*tests. Data are mean s.e.m. **c,** Quantification of *DHFR gene* copy number in MTX resistant colonies from U2OS and Saos-2 cells overexpressing CDT1 and CDC6 compared to non-resistant cells by Real-Time PCR. **d,** Number of MTX resistant colonies upon Cdt1 and/or Cdc6 overexpression, or not, in NIH3T3 cells, corrected to cell survival. NIH3T3 cells, stably overexpressing Cdt1 and NIH3T3 MSCV (stable for empty vector), were transiently transfected with CDC6-Cherry or Cherry as a control and after 48 hours, 10^5^ cells were seeded in 10 cm plates. After 24 hours cells were treated with MTX for 20 days and resistant colonies were fixed, stained and counted; *n*=3 biologically independent replicates, CTRL vs Cdt1 OE, **P< 0.01; CTRL vs Cdt1+Cdc6 OE, **P< 0.01; CTRL vs Cdc6 OE; ns, not significant. Statistical analysis was performed using two-tailed Mann-Whitney *U-*tests. Data are mean ± s.e.m. **e,** *Dhfr* gene copy number quantification by Real-Time PCR in MTX resistant colonies from NIH3T3 cells overexpressing Cdt1 compared to non-resistant cells.

### Multiple DNA damage response mediators are activated following DNA re-replication in cancer cells

We then wished to investigate the cellular responses upon CDT1/CDC6 overexpression in cancer cells. We showed that overexpression of CDT1 and CDC6 in U2OS and Saos-2 cells resulted in overt DNA re-replication, manifested by increased Hoechst integrated density and enlarged nuclei (Fig. 5a, b), elevated DNA replication stress, marked by RPA foci formation (Extended Data Fig. 5a, b) and DSBs, identified by γH2AX (Extended Data Fig. 5c, d). To explore downstream effects of DNA damage evoked by re-replication, we examined activation of double strand break repair mediators. Re-replication was induced in U2OS and Saos-2 cells. Cells were then immunostained for the NHEJ mediators^33^, 53BP1 and RIF1, the HR factor^33^, BRCA1, as well as for RAD52, a recombinase taking part in SSA^34^ and repair of collapsed replication forks through BIR^35^ (Fig. 5c) Image analysis showed activation of all the DSB repair mediators examined in rereplicating cells, revealed by foci formation (Fig. 5 d, e, f, g). Thus, multiple DSB repair pathways are activated following re-replication in cancer cell lines.

**Fig. 5.**
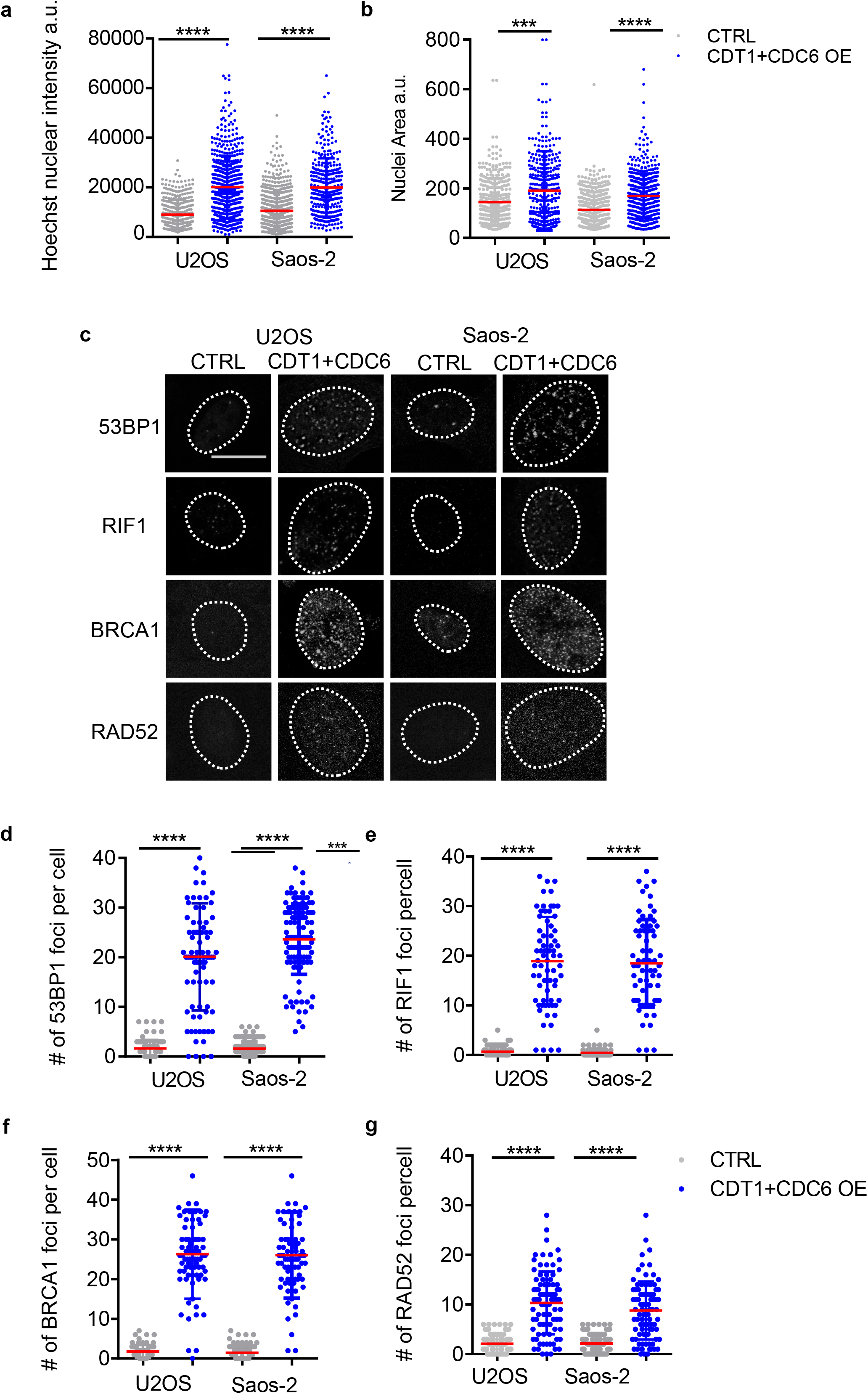
Multiple DNA damage repair mediators are activated upon re-replication in cancer cells. **a,** Quantification of Hoechst integrated density and **b,** quantification of nuclear area in U2OS and Saos-2 cells overexpressing CDT1 and CDC6. U2OS and Saos-2 cells were transiently transfected with CDT1-GFP and CDC6-Cherry or GFP and Cherry, as a control, and after 48 hours the cells were fixed and stained with Hoechst. Scatter plots depict the per cell quantification of nuclear intensity and area where mean ± SD is indicated by red lines. n > 500; CTRL vs CDT1+CDC6 OE, ****P< 0.0001 and ***P<0.001. Statistical analysis was performed using two-tailed Mann-Whitney *U-*tests. **c,** Representative images of U2OS and Saos-2 cells overexpressing CDT1 and CDC6 immunostained for 53BP1, RIF1, BRCA1 and RAD52. Scale bar, 7μm. **d,** Scatter plot depicts the per cell quantification of the number of 53BP1 **e,** RIF1 **f,** BRCA1 and **g,** RAD52 foci from U2OS and Saos-2 cells overexpressing CDT1 and CDC6 where mean ± SD is indicated by red lines; n>100., CTRL vs CDT1+CDC6 OE, ****P< 0.0001. Statistical analysis was performed using two-tailed Mann-Whitney *U-*tests.

### Involvement of ATM and ATR in re-replication induced CNGs

The DNA damage response signaling in mammalian cells is monitored by the central regulators, ATM and ATR^36^. To investigate the possible implication of a checkpoint activation mediated by ATM and ATR in CNGs following re-replication, colony MTX resistance assays were performed in U2OS cells, in the presence or absence of the ATM/ATR inhibitor, caffeine. Inhibition of ATM and ATR decreased the number of MTX resistant colonies in re-replicated cells, suggesting that checkpoint activation following re-replication is required for the formation of CNGs (Extended Data Fig. 6a). Depletion of ATM and ATR did not affect the levels of re-replication (Extended Data Fig. 6c) but weakened the downstream DNA damage response as indicated by reduced H2AX phosphorylation and limited formation of 53BP1 and BRCA1 foci (Extended Data Fig. 6 b, d, e, f). Overall, activation of an intact DNA damage checkpoint is indispensable for formation of CNGs upon re-replication in cancer cells.

### RAD52 participates in the resolution of re-replicating intermediates to generate CNGs

In order to elucidate the repair pathways contributing to CNGs upon re-replication in mammalian cells, we targeted core components of different double strand break repair pathways, such as 53BP1 for NHEJ, BRCA1 and RAD51 for HR and RAD52 for SSA/BIR. The expression of 53BP1, BRCA1, RAD51 and RAD52 was silenced in U2OS cells (Extended Data Fig. 7a, b, c, d), re-replication was induced by CDT1 and CDC6 overexpression and cells were selected with MTX. BRCA1 depletion reduced the number of MTX resistant colonies, while 53BP1 depletion increased their number, suggesting a requirement for a resection-dependent repair mechanism (Fig. 6a). Notably, depletion of RAD52 recombinase, resulted in a pronounced decrease in the number of MTX resistant colonies upon re-replication, while no difference was observed upon RAD51 depletion (Fig. 6a). RAD52 depleted re-replicating cells showed reduced proliferation rate and decreased cell viability (Extended Data Fig. 7e and Fig. 6b) due to enhanced DNA damage response indicated by 53BP1 (Fig. 6c, d) and H2AX (Fig. 6e, f) foci formation. Depletion of other DSB repair components participating in NHEJ (si53BP1) and HR (siBRCA1 and siRAD51) did not cause severe cell death in the presence of re-replication (Extended Data Fig. 7f). Silencing of 53BP1 or BRCA1 resulted in decreased phosphorylation of H2AX in cells overexpressing CDT1 and CDC6, indicating an alternative repair process activated upon re-replication (Extended Data Fig. 7g, h). Moreover, elevated mitotic segregation errors such as anaphase bridges and genomic instability marked by micronuclei formation were also evident in re-replicating cells depleted for RAD52 (Fig. 6g, h, i). Depletion of 53BP1 increased the formation of RAD52 foci indicating that RAD52 and 53BP1 antagonize each other for repair under conditions of re-replication (Extended Data Fig. 7i, j). Overall, RAD52 is important for the resolution of re-replication structures to CNGs.

**Fig. 6:**
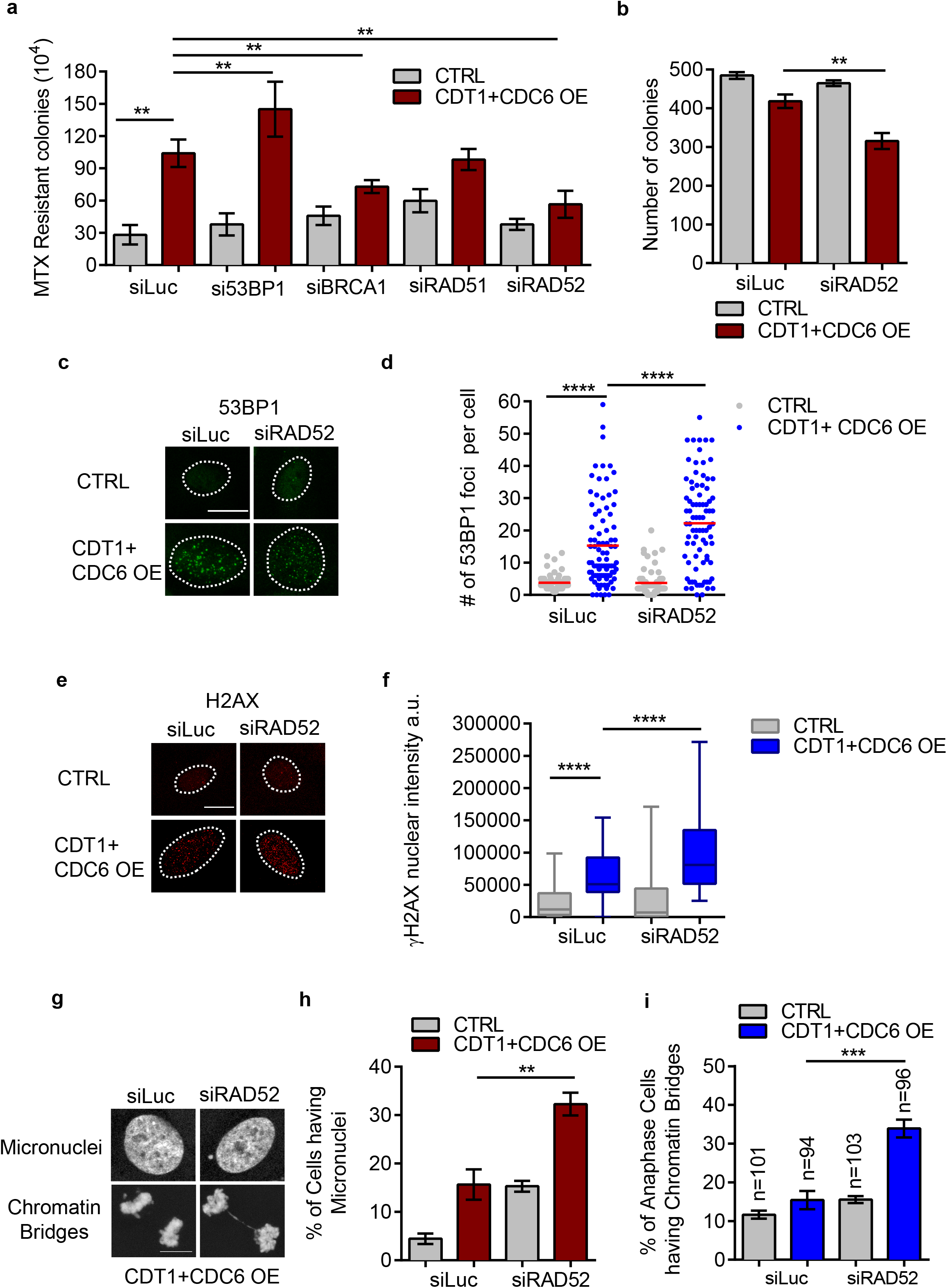
RAD52 participates in CNGs formation following re-replication and its absence in cells overexpressing CDT1 and CDC6 leads to increased genotoxic stress. **a,** Number of MTX resistant colonies upon re-replication induction, or not, in cells depleted, or not, for DSB repair mediators. U2OS cells were depleted for 53BP1, BRCA1, RAD51, or RAD52 and then transfected with CDT1-GFP and Cherry-CDC6 or GFP and Cherry. After 48 hours, 10^4^ cells were plated and treated with MTX for 20 days and resistant colonies were stained and counted. *n*=3 biologically independent replicates, siLuc CTRL vs siLuc CDT1+CDC6 OE, **P< 0.01; siLuc CDT1+CDC6 OE vs si53BP1 CDT1+CDC6 OE, **P< 0.01; siLuc CDT1+CDC6 vs siBRCA1 CDT1+CDC6 OE, **P<0.01; siLuc CDT1+CDC6 OE vs siRAD52 CDT1+CDC6 OE, **P<0.01. Statistical analysis was performed using two-tailed Mann-Whitney *U-*tests. Data are mean ± s.e.m. **b,** Quantification of number of colonies. *n*=3 biologically independent replicates, siLuc CDT1+CDC6 OE vs si RAD52 CDT1+CDC6 OE, **P< 0.01, two-tailed Mann-Whitney *U-*tests. Data are mean s.e.m. **c,** Representative images of U2OS cells immunostained for 53BP1. Scale bar, 7μm. U2OS cells transfected with CDT1-GFP and Cherry-CDC6 and RAD52 RNAi for 48 hours cells were fixed and immunostained. **d,** Scatter plot depicts the per cell number of 53BP1 foci where mean ± SD is indicated by red lines; siLuc CTRL vs siLuc CDT1+CDC6 OE, ****P< 0.0001; siLuc CDT1+CDC6 OE vs siRAD52 CDT1+CDC6 OE, ****P<0.0001.**e,** Representative images of U2OS cells immunostained for γH2AX. Scale bar, 7μm. **f,** Quantification of γH2AX integrated density. The boxes represent the median and quartiles and the whiskers represent the minimum and maximum values; siLuc CTRL vs siLuc CDT1+CDC6 OE, ****P< 0.001; siLuc CDT1+CDC6 OE vs siRAD52 CDT1+CDC6 OE, ****P< 0.0001 For **d,** and **f,** statistical analysis was performed using two-tailed Mann-Whitney *U-* tests **g,** Representative images of U2OS cells having chromatin bridges and micronuclei. Scale bar, 7μm. **h,** Quantification of percentage of cells having micronuclei; siLuc CDT1+CDC6 OE vs siRAD52 CDT1+CDC6 OE, \*\**P*<0.01. **i,** Quantification of percentage of cells having chromatin bridges; siLuc CDT1+CDC6 OE vs siRAD52 CDT1+CDC6 OE. \*\*\**P*<0.001. For **h,** and **i,** *n*=3 biologically independent replicates where performed. Data are mean ± s.e.m. Statistical analysis was performed using two-tailed Student’s t-tests.

### High expression of the licensing factors CDT1 and CDC6 is associated with increased rates of CNGs in cancer patients

To investigate the effect of CDT1 and CDC6 overexpression on CNGs at specific loci in cancer patients, we utilized publicly available data from The Cancer Genome Atlas (TCGA) database^37^. Specifically, the MYC, CCND1, ASPH and ERBB2 loci were examined in breast cancer specimens (Fig. 7a, b, c, d). To determine CNGs at these loci, we used the copy numbers computed by the GISTIC algorithm (Online Methods). For each of the two factors, percentile categorization was used to divide samples in four groups based on their expression of *CDT1* or *CDC6* (Online Methods). In the breast invasive carcinoma dataset, samples with *CDT1* and *CDC6* expression above the upper quartile are significantly correlated with higher percentage of high-level amplification (score=2) in all the loci investigated (Fig. 7a, b, c, d). Similar results were obtained for MYC and CCND1 amplification also in various cancer types (Extended Data Fig. 8a, b, c). We further examined whether the expression of these two licensing factors is linked to increased level of GNGs across the genome. For the cancer datasets analyzed, genes amplified in > 10% of samples were selected and a total amplification score was calculated for each sample as the sum of the GISTIC scores of these genes. In specimens with highest expression of *CDT1* and/or *CDC6*, the total amplification score was significantly higher compared to samples with the lowest expression (Fig. 7e, f, g, h). Taken together, these results suggest that in cancer patients the expression of licensing factors CDT1 and CDC6 are strongly associated with gene amplification.

**Fig. 7.**
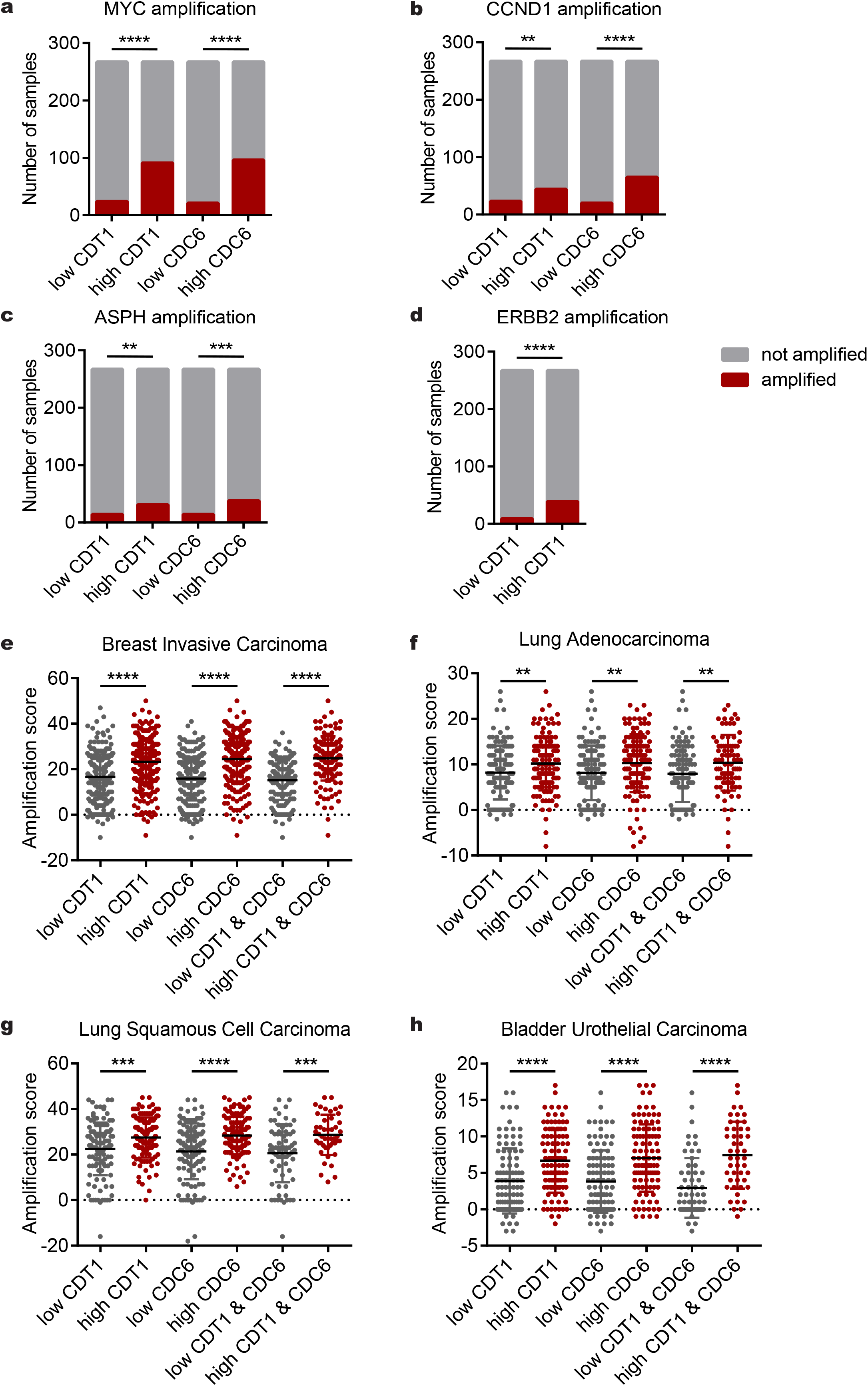
High mRNA levels of the licensing factors CDT1 and CDC6 are highly correlated with increased amplification in specific loci in breast cancer patients and increased overall amplification rate in different cancer types. Patients with breast invasive carcinoma were divided in four groups based on CDT1 or CDC6 expression levels. In each group the number of samples with amplification in **a**, MYC **b**, CCND1 **c**, ASPH and **d**, ERBB2 loci were measured. Statistical significance was calculated with Chi-squared test. To assess the level of gene amplification in total, an amplification score was calculated for every individual patient with **e**, breast invasive carcinoma **f**, lung adenocarcinoma **g**, lung squamous cell carcinoma **h**, bladder urothelial carcinoma. For each cancer dataset, a list of genes that are amplified with frequency higher than 10% and their respective putative copy number (GISTIC score) was obtained. The score was defined as the sum of GISTIC scores of the listed genes. The difference in the amplification scores of the patients with high and low expression of CDT1or CDC6 were compared with Mann Whitney test.

## Discussion

Despite the established importance of CNGs in genome evolution, species evolution and cancer, the mechanisms of their generation remain poorly understood. Through this study we propose re-replication as a driving force for genomic CNGs in different species and different genomic loci. We showed that, re-replication induced by abnormal expression of Cdc18/CDC6 and/ or CDT1 enhances the formation of CNGs in fission yeast, normal and cancer cell lines (Fig. 1a, 4a, b, d). Our data demonstrate that aberrant licensing control can generate CNGs at different genomic loci, affected by local origin activity (Fig. 1d). CNGs are common in cancer^2^, while defective replication licensing has been reported in various tumours^38–41^ and proposed as an initial trigger for malignant transformation^40,42^. Therefore, we provide evidence for a direct link between faulty licensing regulation and CNGs in the cancer genome.

Whole genome sequencing in fission yeast cells selected for re-replication induced CNGs revealed the presence of Mb-long amplicons, mostly present as extrachromosomal repeats, while intrachromosomal inverted structures, which are the most typical BFB-mediated products, were not observed (Fig. 2h, i). Recently, structural analysis of neuroblastoma genomes confirmed the presence of BFB-incompatible amplicons and proposed additional, alternative mechanisms driving CNGs in cancer^43^. Earlier studies in yeasts, have also suggested replication based mechanisms able to drive genome rearrangements under normal conditions^44–47^, while intrachromosomal, tandemly-arrayed amplicons observed in *S. cerevisiae* upon re-replication are also inconsistent with the BFB mechanism^14^. The extrachromosomal nature of re-replication induced CNGs observed in this study for *S. pombe* is in agreement with the site-specific, re-replication driven, extrachromosomal amplicons generated upon overexpression of the KDM4A methyltransferase in mammals^48^.

Genetic analysis, using fission yeast mutants that lack core DNA damage repair components, showed that CNGs at the *sod2* locus require Rad22A and the MMEJ-specific mutant Rad16, but not Rhp51, pointing to the alternative MMEJ pathway for resolution^26,27^ (Fig. 3b, c, e)(Fig. 8). On the other hand, CNGs at the *his5* locus are mediated by Rad51 and hindered by the BIR-specific mutant *cdc27-D1*^28^ (Fig. 3g, h)(Fig. 8). Our results demonstrate that the resolution of re-replication induced amplicons is channeled through homology-mediated mechanisms and that the exact pathways utilized are locus specific and possibly related to the underlying genomic sequence characteristics. This is consistent with findings in *D. melanogaster* showing that distinct repair mechanisms are favored depending on the genomic content to ensure efficient repair and progression of re-replication forks during follicle cells development^49^. The counteractive effect of HR and BIR on *his5* CNGs indicates a competitive relationship between these two pathways. It is possible that HR and BIR antagonize each other for the resolution of re-replication intermediates at this locus and that CNG formation is favored when the classical HR takes over. Antagonistic effects concerning re-replication fork repair has been also reported in *D. melanogaster*^49^. NHEJ does not affect the resolution of CNGs in either loci (Fig. 3a, f), strengthening the argument against a BFB amplification model under re-replication conditions, as BFB necessitates NHEJ during the first step of the sister intra-chromosomal annealing^46,47,50^.

**Fig. 8.**
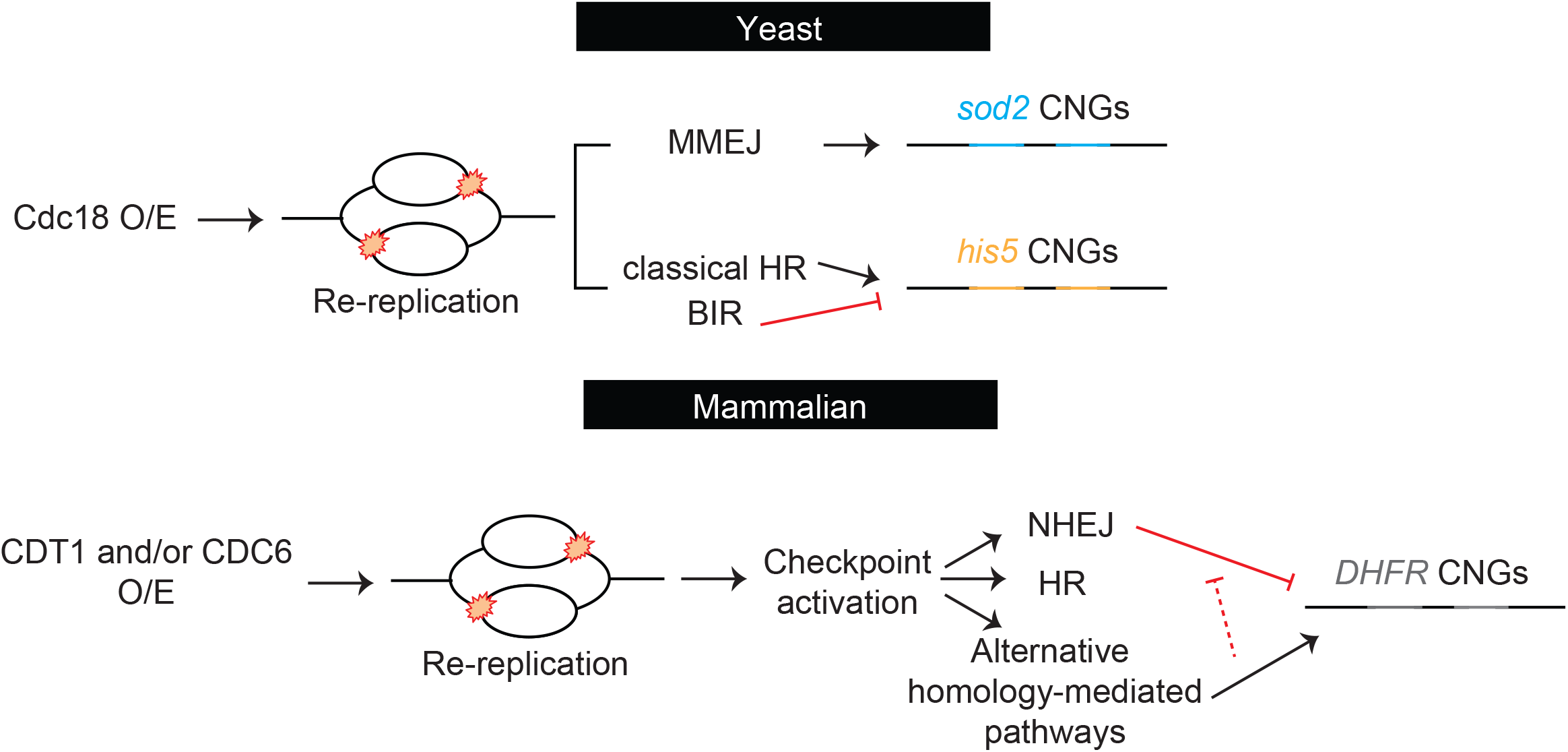
Re-replication induced by aberrant licensing promotes CNG formation across evolution. In fission yeast overexpression of the licensing factor Cdc18 brings about illegitimate origin re-firing and re-replication. Subsequent collapse of the re-replication forks leads to double strand breaks, which in turn elicit the activation of downstream repair pathways, including NHEJ, HR, BIR and the alternative MMEJ. The unstable re-replication intermediates, containing extra copies of specific genes, are resolved through homology-mediated pathways and give rise to stable inheritable CNGs and new phenotypes. Distinct homology-mediated sub-pathways are utilized depending on the genomic locus. Similar to yeast, re-replication induced by ectopic expression of CDT1 and/or CDC6 in humans, leads to fork collapse and double strand breaks. These lesions trigger checkpoint activation and recruitment of different repair mediators. Among them, alternative RAD52-dependent repair pathways favor CNG formation. In contrast, NHEJ suppresses the generation of re-replication induced CNGs and leads to increased genotoxic stress in the absence of RAD52.

Accordingly, pathway analysis in cancer cells shows that ectopic expression of CDT1 and CDC6 leads to increased DNA content, accompanied by replication stress, while multiple DSB repair pathways are activated, including NHEJ, HR and alternative pathways (Fig. 5) (Fig. 8). By silencing key pathway components, we show that the resolution of re-replication intermediates into stable amplicons at the *DHFR* locus is RAD52 dependent (Fig. 6a) (Fig. 8). The dependency on RAD52 points to the error prone repair pathways, BIR or SSA, which have been also recently proposed as the predominant repair routes favored upon p21-driven licensing abnormalities and replication stress^51,52^. RAD52 inactivation not only inhibits CNG formation but also leads to increased DNA damage and mitotic defects, including anaphase bridges and increased lethality (Fig. 6b-i). In contrast, 53BP1 depletion leads to an increase in CNGs (Fig. 6a), suggesting that NHEJ suppresses the formation of re-replication induced CNGs. Notably, RAD52 depletion leads to increased 53BP1 foci, implying a direct competition between the two pathways under re-replication conditions. We suggest that a re-replicating cell could either stabilize its genome by CNG formation through RAD52 dependent pathways and survive or follow alternative repair routes, commonly leading to cell death.

Analysis of TCGA cancer specimens presenting high *CDT1* and *CDC6* mRNA levels revealed increased incidence of CNGs at specific genomic loci, known to contain oncogenes, while the overall amplification rate of the cancer genomes tested was also elevated. Considering that overexpression of licensing factors has been documented in several cancer types^41^ and is able to potentiate malignant transformation^40,42^, these findings point to clinical relevance for re-replication induced CNGs.

CNGs enhance the adaptive potential of cancer cells^53,54^, facilitating clonal expansion, tumor heterogeneity, drug resistance, and tumor aggressiveness^4,54,55^ and therefore understanding the mechanisms of their generation is critical. Herein, we highlight re-replication caused by perturbed replication licensing, as an evolutionary-conserved, driving mechanism for the formation of CNGs, with applications in cancer. We provide insight into the underlying molecular mechanisms, which can be exploited to uncover potential therapeutic targets in order to obstruct the generation of CNGs and their subsequent effects in *CDT1* and *CDC6* overexpressing tumor cohorts.

Several recent studies propose that specific mutational signatures in cancer are accompanied by distinct structural variants^48,56–60^. *CDT1* and *CDC6* overexpression could act as a mutator phenotype characterized by increased CNGs, which in turn could be used as a classification criterion for cancer subtypes.

## Online Methods

### Yeast Strains

The genotypes of all the strains used in this study are shown in Supplementary Table 1. All the strains constructed were derived by the parental strain ZP151(*d55P6-cdc18*)^61^. *d55P6-cdc18* lacks 149 amino acids in the N terminus, including five out of six cdc2-targeted phosphorylation sites, and additionally bears a threonine to alanine substitution at amino-acid 374 that disrupts the sixth phosphorylation site^29^. For the construction of the strains bearing the *his5* gene next to ars3002 or ars727 the following procedure was followed: the *ura4* gene was amplified from the plasmid pura4script^62^ with oligos 553,554 and targeted to the endogenous *his5* locus of the ZP151 strain by the standard PCR-based method^63^ giving rise to ZP376 which is deleted for the endogenous *his5* gene. Two synthetic constructs were used: p918 include the target sequence next to origin ars3002 between SalI and BamHI restriction sites and the *his5* gene with its regulatory elements between BamHI and XmaI restriction sites and p919 carried the second target sequence next to ars727 between SalI and BamHI restriction sites. The SalI-XmaI fragment of p918 sequence was cloned in the respective sites of the pFA6A-kanMx6 plasmid^63^ and the final construct p920 was transformed in ZP376 and targeted to ars3002 (ZP408). Next, the SalI-BamHI fragment of p920 was substituted by the second target sequence from plasmid p919, transformed in ZP376 and targeted to ars727 (ZP466). The sites of integration were validated by colony PCR using oligos 628-631, 626-630 and 679-680, 681-682 for ars3002 and ars727, respectively. Plasmids carrying two or three copies of the *his5* gene were constructed as follow: the BamHI-BglII fragment from p918 was cloned in the BglII site of p920. The number of copies and their orientation were checked by restriction digests. DDR mutant strains *pku70Δ*, *rad22AΔ*, *rad51Δ*, *rad16Δ* were kindly provided by the Nurse lab, while the *cdc27-D1* was provided by Timothy Humphrey’s lab. These mutants were crossed in ZP408 and the resulting strains were used for the colony formation assays. Deletion of each DNA damage repair component was verified by colony PCR. A list of the oligos used are available is Supplementary Table 2.

### Colony formation assay in yeast

Strains in a *nmt1-d55P6-cdc18* background were cultured overnight in EMM supplemented with appropriate amino acids and 5 μgr/ml thiamine (T). The cells were diluted in EMM+T and grown until mid-exponential phase. Thiamine was removed by three washes in EMM. The cells were appropriately diluted in EMM or EMM+T as a control and cultured for specific hours to reach the desired induction time. The exact number of cells was counted, and both cells were plated in EMM+T to estimate the survival rate and in EMM+T+40mM LiCl (Sigma Aldrich-62476) to select for *sod2* amplification or EMM+T+5 mM 3AT (Sigma Aldrich-A8056) for the *his5* amplification. For the DNA damage repair mutant analysis *nmt1-d55P6-cdc18* wild type and mutant strains were processed in parallel following the steps mentioned before. 3AT resistance assays were not performed for the Rad22AΔ mutant because of technical issues. 3AT selection requires absence of histidine in the culture medium and therefore the assays should be performed in EMM-his+T. However, for undefined reasons the Rad22AΔ mutant presented difficulties growing in minimal medium. For the Rad22AΔ mutant, LiCl selection was performed in YES+40 mM LiCl+T.

### Spot assay

The exact cell number of mid exponential growing cultures was counted using a Neubauer cell counting chamber. 10μl of serial dilutions (10^8^-10^3^ cells/ml) were spotted on plates containing 5, 10, 15, 20mM 3-AT. Plates were incubated for 3 days at 32°C. The 3-AT concentration that was used for the *his5* colony formation assays was titrated using strains that carry artificially two or three copies of the *his5* gene with its regulatory elements next to each ORI (Extended Data Fig. 1d, e). 5 mM of 3-AT was chosen as the appropriate concentration to distinguish between single and multiple copies of the gene.

### Yeast genomic preparation

The protocol for small scale genomic preparation was followed. 10ml of cells in stationary phase were resuspended in 1 ml 50 mM citrate/phosphate pH 5.6, 40 mM EDTA, 1.2 M sorbitol. 0.5 mg Zymolyase 100T (nacalai tesque-07665-55) were added for cell wall digestion. Spheroplasts were spinned down and lysed in 1% SDS at 65°C for 1 hour. 175 μl of 5 M potassium acetate was added, the samples were kept on ice for 5 min and centrifuged. 0.5 ml of the supernatant was precipitated using isopropanol, the pellet was washed with 70% ethanol, resuspended in TE buffer containing 50 μg/ml RNAase and incubated in 65°C for 10 minutes. Genomic DNA isolation was performed using phenol/chloroform extraction and the resulting DNA was resuspended in 30 μl TE buffer. For genomic preparation for WGS the same steps were followed until the cell wall digestion. After that the spheroplasts were resuspended in 500 μl spooling buffer (75 mM NaCl, 25 mM EDTA, 1% SDS) containing 100 μg/ml Proteinase K. The samples were incubated overnight at 55°C with gentle agitation. 50 μg/ml RNAase were added, and the samples were incubated at 37°C for 30 minutes. 125 μl of saturated NaCl and 625 μl of isopropanol were added. The samples were spinned down for 10 minutes at 13000 rpm. The pellet was washed with 70% ethanol and air dried. Reconstitution of the pellet was achieved in 10 mM Tris pH 8.0 at 55°C with gentle agitation.

### Whole genome sequencing – Structural analysis

Library preparation and whole genome sequencing was performed at the Genecore facility in EMBL. Paired end sequencing, at 80x depth was performed using the Illumina HiSeq2500 sequencer. Four LiCl resistant, four 3-AT resistant and two non-resistant control colonies were sequenced. The number of total reads per 1000 base pairs was normalized to the control and the copy number profile was plotted for each resistant sample. To assess possible technical variations of the sequencing method, two independent genomic isolations of the same 3-AT colony were prepared and sequenced (3AT 1.1, 3AT 1.2). The copy number profiles between these two samples were identical. The structural analysis was carried out manually by assessing parameters like pair read orientation, insert size and chromosome of paired reads at the amplicon boundaries.

### Pulse Field Gel Electrophoresis and Southern Blot

Agarose plugs were prepared from 150 ml mid exponential phase cultures. The cells were washed twice with CSE buffer (20 mM citrate/phosphate pH 5.6, 40 mM EDTA, 1.2 M Sorbitol), resuspended in 10 ml CSE with 3 mg Zymolyase-100T and digested for 1 hour at 37°C. The cells were pelleted and resuspended at 6×10^8^ cells/ml in TSE (10 mM Tris HCl pH 7.5, 45 mM EDTA, 0.9 M Sorbitol) and the same volume of 1% low melting agarose was added. The samples were immediately dispensed in plug molds. The plugs were incubated in 0.25 M EDTA, 50 mM Tris HCl pH 7.5, 1% SDS for 90 minutes at 55°C to lyse the cells. The plugs were transferred to 1% lauryl sarcosine, 0.5 M EDTA pH 9.5,0.5 mg/ml Proteinase K and digest at 55°C for 48h. Fresh Proteinase K was added after 24 hours. Plugs were stored at 4°C.

For analysis of whole chromosomes, plugs were loaded in 0.8% pulsed-field certified grade agarose gels (Biorad) and electrophoresed in 1x TAE at 14°C in a CHEF-DR III pulsed-field gel apparatus (Biorad) at 2 V/cm for 72 h, using one block. The parameters were: initial switch time 20 minutes-final switch time 60 minutes (linear fashion), 106° included angle. For the separation of smaller fragments plugs were loaded in 1% gel and electrophoresed in 0.5x TBE at 14°C for 24 hours using one block with the following parameters: initial switch time 28.6 seconds-final switch time 228 seconds (linear fashion), 120° included angle, 6 V/cm. Gels were stained with 0.5 μg/ml ethidium bromide in electrophoresis buffer.

For the Southern Blot DNA was transferred in a positively charged nylon membrane (Amersham-RPN2020B) following standard procedures. Hybridizations were carried out at 68°C with ~2×10^6^ c.p.m. of randomly primed probe. After washing, the membranes were exposed to autoradiography films.

### Probing

The desired fragments were amplified from the genome using colony PCR and purified by gel extraction. The extracted fragments were labelled using the Prime it II Random Primer Labelling kit (Agilent-300385) and the unincorporated nucleotides were removed using the ProbeQuant G-50 Micro columns (GE Healthcare-28903408) according to manufacturer’s instructions, respectively.

### Imaging and cell length analysis in yeast

Cells were fixed with 4% paraformaldehyde (Sigma Aldrich-P6148), mounted onto glass slides with Mowiol 4-88 and brightfield images were acquired using a Leica SP5 confocal microscope (63x lens). Cell length values were measured for 100 cells per condition using the PointPicker plug-in of ImageJ/FIJI analysis software.

### Flow cytometry

For yeast cells 1 ml of mid-exponential growing cells (O.D. 0.5) was centrifuged at 2000 rpm for 5 minutes, washed once with water, fixed in 1 ml ice cold ethanol and stored at 4°C. Before processing, 300 μl of cells were pelleted, ethanol was discarded, and the cells were rehydrated in 1 ml 50 mM sodium citrate. Cells were again pelleted, resuspended in 0.5 ml 50 mM sodium citrate containing 10 mg/ml RNAase (Sigma Aldrich-R5125) and incubated at 37°C for at least 2 hours. 0.5 ml of sodium citrate containing 4 μg/ml Propidium Iodine (Sigma Aldrich-P4864) was added to stain DNA, the samples were sonicated and immediately processed using a BD FACSCalibur flow cytometer. Data were collected and analyzed using FlowJo.

NIH3T3 MSCV and NIH3T3 *Cdt1* cells were harvested by trypsinization and fixed using ice-cold 70% ethanol (added drop wise) and stored at −20°C overnight. Then cells were centrifuged at 2000g for 15 min at 4 °C, rinsed twice in 1X PBS and then stained with 2 mg/ml propidium iodide (Sigma Aldrich-P4864) and 100 g/ml RNase A (Sigma Aldrich-R5125) dissolved in 1X PBS for 30 min at 37°C. Cellular DNA content was assessed on a FACS Calibur flow cytometer (BD Biosciences) and analyzed with Flowjo.

### Cell culture

Human U2OS (ATCC) and Saos-2 parental cells (from V. Gorgoulis), were maintained in Dulbecco’s modified Eagle’s medium (Invitrogen, Cat. No. 41966-029), supplemented with 10% fetal bovine serum (FBS; Invitrogen, Cat. No. 10270-106) and antibiotics penicillin/streptomycin (Invitrogen, Cat. No. 15140-122) at 37°C in a humid atmosphere with 5% CO2. NIH3T3 MSCV and NIH3T3 *Cdt1* stable cells lines (from K. Choi) were maintained in Dulbecco’s modified Eagle’s medium, supplemented with 10% fetal bovine serum and antibiotics penicillin/streptomycin and puromycin 1μg/ml (Invitrogen, Cat. No. A11138-03). Cells were routinely subjected to mycoplasma testing and found to be negative.

### Plasmid DNA and siRNA Transfections

For plasmid transfections, cells were transfected at 70-90% confluency prior to sample collection with the indicated plasmids using Lipofectamin 2000 transfection reagent (ThermoFisher Cat. No. 11668027). Plasmids used: GFP and CDT1-GFP^64^, mCherry-CDC6, Myc-CDC6. For knockdown experiments, cells were transfected at 70-90% confluency prior to sample collection with the indicated siRNA using Lipofectamine RNAiMax transfection reagent (ThermoFisher Cat. No. 13778075) according to manufacturer’s instructions. siRNA sequences used: siLuciferase (48h, 80nM, 5’ CGUACGCGGAAUACUUCCAdTdT 3’), si53BP1 (48h, 80nM, 5’ GAGAGCAGAUGAUCCUUUAdTdT 3’), siBRCA1 (48h, 80 nM, 5’ CAGCUACCCUUCCAUCAUAUUdTdT3’), siRAD51 (48h, 80 nM, 5’ GGGAAUUAGUGAAGCCAAAdTdT 3’), siRAD52 (48h, 80nM, 5’ GGUCCAUGCCUUUAAUGUUdTdT 3’). All siRNAs were purchased from Eurofins MWG. Efficiency of siRNAs is depicted in Extended Data Fig. 6.

### Cell growth, survival and colony formation assays in mammalian cells

For cell growth assay 10^5^ cells were seeded per well in 6-well plates. After 24 hours, cells were transfected with indicated siRNAs using RNAimax (Invitrogen), followed by indicated plasmid transfection using Lipofectamine 2000 (Ιnvitrogen). After 24,48 and 72 hours cells were trypsinized and counted. For cell survival and colony formation assays, following any treatment conditions, equal number of cells (500 cells for cell survival and 10^4^ or 10^5^ for drug resistance) were seeded onto 10 cm culture dishes, maintained at 37 °C in an atmosphere of 5% CO_2_. Cells were allowed to form colonies for 10 days, for cell survival assays or for 20 days for drug resistance assays. Colonies were stained using 0.4% methylene blue in 50% methanol and quantified using ImageJ/FIJI analysis software.

### MTT assay

Cell viability was determined using the 3-(4,5-dimethylthiazol-2-yl)-2,5-diphenyltetrazolium bromide (MTT) assay. For this, 3×10^4^ cells per well were plated onto 24-well plates. After treatment 10% volume of 5 mM MTT (Applichem Cat. No. A2231) solution in PBS was added to wells and incubated for 3 h at 37 °C in a 5% CO_2_ atmosphere. The metabolized insoluble formazan crystals were then solubilized using DMSO and the absorbance was measured at 492 nm on a MR-96 A photometer (Mindray, Shenzhen, China).

### Immunofluorescence

Cells were seeded on autoclaved glass poly-D-lysine (Sigma-P1399) coated coverslips and after treatment pre-extracted with ice-cold PBS containing 0.2% Triton X-100 for 2 min on ice before fixation with 4% paraformaldehyde for 10 min. Cells were blocked with PBS containing 10% FBS, 3% BSA for 1hr and then were incubated with primary antibodies overnight at 4°C. Primary antibodies used: 53BP1 (1-1000, Abcam Cat. No. ab21083), RIF1 (1-500, A300-569A, Bethyl Laboratories), BRCA1 (1-1000, D-9, sc-6954, Santa Cruz), Phospho-Histone H2A.X-Ser139 (1-1000, 05-636, Merk Millipore), RAD52 (1-100, F-7, sc-365341, Santa Cruz Biotechnology). Following primary antibody incubation, cells were incubated for 1 hour with secondary antibodies conjugated with Alexa Fluor 488 (1-1000, Invitrogen), Alexa Fluor 568 (1-1000, Invitrogen), Alexa Fluor 647 (1-1000, Invitrogen). Cell nuclei were counterstained with Hoechst 33342 (Invitrogen, Cat. No. H3570) and mounted with Mowiol 4-88 (Calbiochem) on glass slides. Images were recorded in a Leica TCS SP5 confocal laser scanning microscope (Leica Microsystems) using a 63X oil immersion objective or in a Nikon Eclipse TE2000-U using a 40X air immersion objective. For micronuclei and anaphase bridges detection, cells were stained following indicated treatment with Hoechst 33342 (Invitrogen, Cat. No. H3570) and micronuclei and anaphase bridges were scored with ImageJ/FIJI analysis software. For quantification of Hoechst integrated density, mean intensity and number of foci, custom ImageJ/FIJI macros were used.

### Immunoblotting

Following any indicated treatement, cells were collected using cell scraper and total cell lysates were prepared by lysing cell pellets immediately in SDS-PAGE loading buffer (160 mM Tris-HCl pH 6.8, 4% SDS, 20% glycerol, 100 mM DTT, and 0.01% bromophenol blue). The lysates were incubated at 95% for 5 min and stored at −80 °C. Equal amounts of proteins were analysed on 10% acrylamide – 0.1% bisacrylamide gel for 1hr, (80V, room temperature) and blotted for 90 min (42V, room temperature) on PVDF membrane (Millipore, IPVH00010). Membranes were blocked in 5% milk in 0.1% PBS-Tween 20 for 30min and incubated in 2.5% milk in 0.1% PBS-Tween 20 with primary antibodies overnight. Primary antibodies used; RAD51 (1-500, Bio Academia Cat. No. 70-001); RAD52 (1:500, F-7, sc-365341, Santa Cruz Biotechnology); a tubulin (1-20000, Sigma Aldrich Cat. No. Τ5168). Membranes were incubated with HRP-linked secondary antibodies (Santa Cruz Biotechnology) for 1h at room temperature (in blocking buffer). Membranes were washed three times with 0.1% PBS-Tween 20, after primary and secondary antibody incubations and the signal was developed using ECL detection reagent (Thermo Scientific Cat. No. 32106).

### Quantitative real-time PCR

Genomic DNA from yeast cells was prepared as described above. For mammalian cells genomic DNA was extracted using the Nucleospin Genomic DNA from tissue kit (Cat. No. 740952). Genomic DNA concentration and purity was measured using Nanodrop. Gene copy number was assessed by quantitative real-time PCR (Applied Biosystems StepOne), using the Kapa SYBR Fast qPCR kit (Kapa Biosystems, KK4605). The copy number of the Human *GAPDH* and mouse *HPRT* or the ars727 genomic region were used for normalization in mammalian and yeast cells, respectively. 8 ng/μl of genomic DNA was used per reaction in both yeast and mammalian cells. For qPCR data analysis, the REST-MCS beta software was used. The sequence of the oligos used are available in Supplementary Table 2.

### Chemicals

The following reagents were used to treat the cells for the indicated time at the indicated final concentrations before sample collection: Lithium-Chloride-LiCl (Sigma Aldrich-62476), 3-Amino-1,2,4-triazole 3AT (Sigma Aldrich-A8056), Methotrexate-MTX (Sigma Aldrich; Cat. No. M8407), MTT Thiazolyl blue tetrazolium bromide (Applichem Cat. No. A2231), Caffeine (Sigma Aldrich Cat. No. C0750)

### Statistical analysis

Five independent replicates of each colony formation assay in fission yeast were performed and the resulting values were analyzed using the unpaired, non-parametric, two-tailed Mann-Whitney test for each condition compared. Mann-Whitney and Students t-test was also used to statistically assess differences in experiments involving human and mouse cell lines. Statistical analysis was performed using GraphPad Prism 6 (GraphPad Software).

### TCGA data analysis

High throughput genomic data for the different cancer types analyzed in this study, were obtained from cBio Cancer Genomics Portal^65,66^ using cgdsr package in R (https://github.com/cBioPortal/cgdsr). For each cancer type samples with both expression and CNA data were obtained. Putative copy number for each gene was determined through copy number analysis with GISTIC algorithm (Genomic Identification of Significant Targets in Cancer)^67^. A GISTIC-score was assigned to each gene based on their copy number level. A score=−2 corresponds to deep (homozygous) deletion, −1 to heterozygous deletion, 0 is used to describe the diploid situation and scores 1 and 2 corresponds to low-level gain and high-level amplification respectively. For the estimation of the relative expression levels of the licensing factors *CDT1* and *CDC6*, mRNA expression (RNA Seq V2 RSEM) data were used. The Cancer datasets used, and the respective cancer study ID are the following:

| Cancer Study ID | Cancer Type | Number of samples |
| --- | --- | --- |
| blca_tcga | Bladder Urothelial Carcinoma (TCGA, Provisional) | 404 |
| brca_tcga | Breast Invasive Carcinoma (TCGA, Provisional) | 1068 |
| luad_tcga | Lung Adenocarcinoma (TCGA, Provisional) | 512 |
| lusc_tcga | Lung Squamous Cell Carcinoma (TCGA, Provisional) | 496 |

For every data set, samples were divided in quartiles based on the mRNA expression of *CDT1* or *CDC6*. Samples in the first quartile of the *CDT1* or *CDC6* distribution were considered as low *CDT1* or *CDC6* samples while samples in the fourth quartile were considered as high *CDT1* or *CDC6* samples. As amplified genes were considered those with GISTIC score=2, while genes with GISTIC score<2 were considered not amplified.

To assess the level of amplification across the entire genome a new metric called amplification score was employed. For each cancer dataset used in this study, a list of genes that are amplified with frequency higher than 10% and their respective putative copy number (GISTIC score) was obtained. In cases where multiple genes were located in the same chromosomal coordinates only one representative gene was used for the analysis. The amplification score for every sample was defined as the sum of the GISTIC scores of the selected gene set. The gene sets that were taken into account to calculate amplification score in each cancer type are represented in Table S3.

## Acknowledgments

We thank Dr. Nickolaos Nikiforos Giakoumakis for assistance with image analysis and Dr. Alexandra Kanellou for the construction of the mCherry-CDC6 plasmid. We thank the Advanced Light Microscopy Facility of the University of Patras. This work was supported by the European Research Council [ERC-StG 281851 and ERC-PoC 755284]; by the State Scholarships Foundation of Greece [PhD fellowship to P.N., S.M. and E.K.] and by the Hellenic Foundation for Research and Innovation (HFRI) [PhD fellowship to M.P.]. The study was also supported by the project “Bioimaging-GR” (MIS 5002755), implemented under the Action “Reinforcement of the Research and Innovation Infrastructure”, funded by the Operational Programme “Competitiveness, Entrepreneurship and Innovation” (NSRF 2014-2020) and co-financed by Greece and the EU. Funding for open access charge: Bioimaging-GR.

## Author’s Contribution

P.N. designed, performed and analysed experiments in fission yeast and wrote the manuscript with contribution from all authors. M.P. designed, performed and analysed experiments in mammalian cells. S.M. performed TCGA data analysis. E.K. performed experiments with checkpoint mutants in fission yeast. I.E.S initiated studies of drug resistance in mammalian cells, O.P. assisted with mutant analysis in fission yeast. I.S.S. provided equipment and expertise for PFGE. V.B. performed WGS. S.T. co-supervised the study. Z.L. conceived and supervised the study and finalized the manuscript. All authors read, corrected and approved the manuscript.

## Data Availability

The Whole Genome Sequencing raw data files have been deposited in the Sequencing Read Archive-SRA with the identifier PRJNA625927. All data supporting the findings of this study are available from the corresponding author on reasonable request. Source data for Extended Data Fig. 7 are presented with the paper.

---

**Extended Data Fig. 1. a**, Multiple cell numbers for control and re-replicating cultures (23 hours induction) were plated in non-selective medium and the survival in both conditions was assessed and plotted as percentage. **b**, The relative copy number of the *sod2* locus in LiCl resistant colonies arising under normal conditions was assessed by qRT PCR. **c**, Re-replication was induced for different time intervals and the number of LiCl resistant colonies under normal or re-replicating conditions was counted for each timepoint. The number of resistant colonies in the re-replication condition was normalized to the control in each case and plotted. **d**, Strains carrying the *his5* gene in one, two or three copies next to ars3002 (top) or ars727 (bottom) were used to determine the appropriate 3AT concentration for the respective colony resistance assays. The genotypes of the constructed strains are presented. **e**, **f**, The copy number of the *his5* gene in each strain was validated by qRT-PCR for ars3002 and ars727, respectively **g**, The strains were cultured in the presence of thiamine and serial dilutions from each culture were spotted on medium containing increasing concentration of 3-AT both for ars3002 and ars727. The parental strain was also included. 5mM of 3-AT were sufficient to select for multiple versus single copies of the *his5* locus in both genomic locations. **h**, **i**, Determination of the relative copy number of the *his5* locus next to ars3002 (**h)** or **(i)** ars727 in 3-AT resistant colonies grown under normal conditions by qRT PCR.

**Extended Data Fig. 2. a**, **b**, Genome-wide copy number profiles for LiCl (**a**) and 3-AT (**b**) resistant colonies. Total reads per 1000 base pairs in each resistant colony were normalized to the respective total reads of the non-resistant control and plotted. γH2A enriched sites indicating Common Fragile Sites (CFS) are depicted as yellow circles. **c**, Full chromosome analysis of 3AT and LiCl resistant colonies by PFGE. 3AT and LiCl resistant colonies were cultured in the presence of thiamine, under 5 mM 3AT and 20 mM LiCl selection, respectively. Genomic DNA was isolated in agarose plugs for control, 9 3AT and 10 LiCl resistant colonies. Lanes 1, 23: *S. cerevisiae* chromosomal DNA used as size marker (m). Lanes 2,12: non-re-replicating, non-resistant control strains (c), lanes 3-11: 3AT resistant colonies, lanes 13-22: LiCl resistant colonies. **d**, LiCl samples were analyzed by PFGE using low molecular weight electrophoretic conditions to determine the exact size of the extra bands representing the extrachromosomal amplicons. **e**, Full chromosome PFGE followed by Southern blot was performed for 3AT resistant colonies using a probe against the ectopically integrated *his5* locus at Chr.III.

**Extended Data Fig. 3. a-e**, FACS analysis in wild type cells and DNA damage repair mutants. Cells were cultured in the absence of thiamine and FACS samples were collected at several timepoints upon re-replication induction. Samples were stained for PI to monitor DNA content. Timepoints with comparable re-replication levels between the wild type and each mutant were selected for the performance of the respective colony formation assays. The selected timepoints are presented as overlaid histograms. **f**, Schematic representation of the wild type and the N-terminally truncated version of Cdc18 (*d55P6-cdc18*). **g**, FACS analysis for the *nmt1-d55P6-cdc18* strain. Re-replication was induced for 21 hours in wt and *rad3Δ* cells. The DNA content was monitored by PI staining and the respective profiles are presented. **h**, Cell length analysis in cell strains overexpressing *d55P6-cdc18*, in the presence (wt) or absence (*rad3Δ*) of Rad3. Re-replication was induced for 21 hours, or not, and cells were fixed with PFA. Microscopy pictures were taken and the cell length of 100 cells per specimen was measured using ImageJ. In the absence of Rad3 the *d55P6-cdc18* strain is not able to arrest the cell cycle upon re-replication induction, indicated by the absence of the typical elongated phenotype. **i**, Re-replication was induced or not, in wild type and *rad3Δ nmt1-d55P6-cdc18* cells. 2×10^8^ cells were plated in the presence of 40mM LiCl under non-re-replicating conditions and the number of LiCl resistant colonies upon re-replication, or not, corrected to survival, is presented. **j**, The number of colonies in the re-replicating condition for both strains is also presented as fold change over the control, non-re-replicating condition. The non-parametric Mann-Whitney statistical test was used to assess the significance of differences in each case.

**Extended Data Fig. 4. a,** IC50 survival curves of U2OS and Saos-2 treated with different concentrations of MTX determined by MTT. **b,** Quantification of the number of colonies from U2OS and Saos-2 cells overexpressing CDT1 and CDC6 compared to control cells. 500 U2OS and Saos-2 parental cells transiently transfected with CDT1-GFP and CDC6-Cherry for 48hrs were seeded in 10cm plates in normal medium for 10 days to determine cell survival. Colonies were fixed, stained and counted. **c,** *DHFR* copy number quantification in MTX resistant U2OS and Saos-2 compared to non-resistant cells, using Real-Time PCR. **d,** IC50 survival curves of NIH3T3 MSCV (control) and NIH3T3 Cdt1 (Cdt1 OE) cells treated with different concentrations of MTX determined by MTT. **e,** Quantification of the number of colonies from NIH3T3 overexpressing Cdt1 and/or Cdc6. 500 NIH3T3 MSCV and NIH3T3 Cdt1 cells, transiently transfected for 48 hours with Cherry-CDC6 were seeded in 10cm plates in normal medium for 10 days to determine cell survival. Colonies were fixed, stained and counted. **f,** *Dhfr* copy number quantification in MTX resistant NIH3T3 cells to non-resistant cells, using Real-Time PCR (NE: Normal Expression). **g,** Flow cytometry profiles of NIH3T3 MSCV and NIH3T3 Cdt1 cells. DNA stained with PI. **h,** Representative images of NIH3T3 control and NIH3T3 Cdt1 OE cells immunostained for γH2AX. NE: Normal Expression, OE: Overexpression. **i,** Scatter plot depicting per cell γH2AX integrated density. Mean ± SD is indicated by red lines. n > 100; Cdt1 NE vs Cdt1 OE, ****P< 0.001. Statistical analysis was performed using two-tailed Mann-Whitney *U-*tests.

**Extended Data Fig. 5. a,** Representative images of U2OS cells overexpressing CDT1 and CDC6 immunostained for RPA. U2OS cells were transfected with CDT1-GFP and Cherry-CDC6 and after 48 hours cells were fixed and immunostained. Cell nuclei counterstained with Hoechst. **b,** Quantification of the percentage of cells with RPA foci. **c,** Representative images of U2OS cells overexpressing CDT1 and CDC6 immunostained for γH2AX. Cell nuclei counterstained with Hoechst. d, Quantification of the percentage of cells with γH2AX foci. For **b, d,** CTRL vs CDT1+CDC6 OE, \*\*\*\**P<*0.0001, *n*=3 biologically independent. Data are mean ± s.e.m. Statistical analysis was performed using two-tailed Student’s t-tests.

**Extended Data Fig. 6. a,** Number of MTX resistant colonies of U2OS cells overexpressing CDT1 and CDC6 or not, in the presence or the absence of caffeine. U2OS cells were transiently transfected with CDT1-GFP and Cherry-CDC6 or GFP and Cherry, as a control, and treated or not with caffeine and 48 hours later 10^4^ cells were seeded in 10 cm cell culture plates. Cells were treated with MTX for 20 days and then resistant colonies were fixed, stained and counted. *n=*3 biologically independent replicates; – caffeine CTRL vs ‒caffeine CDT1+CDC6 OE, **P< 0.01; -caffeine CDT1+CDC6 OE vs +caffeine CDT1+CDC6 OE, \**P*<0.05. Statistical analysis was performed using two-tailed Mann-Whitney *U-*tests. Data are mean s.e.m **b,** Representative images of cells overexpressing CDT1 and CDC6 and treated with caffeine immunostained for 53BP1, BRCA1 and γH2AX. Scale bar, 7μm. U2OS cells were transiently transfected with CDT1GFP and CDC6-Cherry or GFP and Cherry, as a control, in the presence or absence of caffeine. After 48 hours cells were fixed and immunostained. Nuclei counterstained with Hoechst. **c,** Scatter plot depicts per cell Hoechst integrated density in U2OS cells overexpressing CDT1 and CDC6 and treated with caffeine; -caffeine CTRL vs ‒caffeine CDT1+CDC6 OE, \*\*\*\**P*<0.0001; ns: not significant. **d,** Scatter plots depict per cell number of 53BP1 and **e,** BRCA1 foci in U2OS cells overexpressing CDT1 and CDC6 treated with caffeine. F. Scatter plots depict per cell γH2AX integrated density of U2OS cells treated with caffeine and overexpressing CDT1 and CDC6;-caffeine CTRL vs ‒caffeine CDT1+CDC6 OE, \*\*\*\**P*<0.0001; -caffeine CDT1+CDC6 OE, \*\*\*\**P*<0.0001. Statistical analysis was performed using two-tailed Mann-Whitney *U-*tests.

**Extended Data Fig. 7 a,** Efficiency of siRNA-mediated depletion of 53BP1 and **b,** BRCA1 by immunofluorescence in U2OS cells Scater plots depict the per cell integrated density of 53BP1 and BRCA1. siLuc vs si53BP1, ****P<0.0001; siLuc vs siBRCA1, ****P<0.0001. Statistical analysis was performed using two-tailed Mann-Whitney *U-*tests. **c,** Western blot analysis of U2OS treated with siRAD51 and **d,** siRAD52. a-Tubulin was used as loading control. Uncropped immunoblots are provided in source data. **e,** Cell proliferation assay of U2OS cells overexpressing CDT1 and CDC6 and depleted for RAD52. The number of cells was determined 24, 48 and 72 hours post transfection. **f,** Number of surviving colonies from U2OS cells overexpressing CDT1 and CDC6 and depleted for either 53BP1, BRCA1, RAD51. U2OS cells were transfected with siRNAs targeting 53BP1, BRCA1 or RAD51 and expressing vectors CDT1-GFP and CDC6-Cherry. After 48 hours 500 cells were seeded from each condition in 10cm culture plates and colonies were fixed and counted after 10 days. **g,** Representative images of U2OS cells overepxressing CDT1 and CDC6 and depleted for 53BP1 or BRCA1 immunostained for γH2AX. Scale bar, 7μm. **h,** Quantification of γH2AX integrated density. The boxes represent the median and quartiles and whiskers represent the minimum and maximum values; siLuc CTRL vs siLuc CDT1+CDC6 OE, ****P< 0.0001; siLuc CDT1+CDC6 OE vs si53BP1 CDT1+CDC6 OE, ****P< 0.0001; siLuc CDT1+CDC6 OE vs siBRCA1 CDT1+CDC6 OE, ****P<0.0001. Statistical analysis was performed using two-tailed Mann-Whitney *U-*tests. **i,** Representative images of U2OS cells overexpressing CDT1 and CDC6 and depleted for 53BP1, immunostained for RAD52. Scale bar,7μm. **j,** Scater plot depicts the per cell number of RAD52 foci in U2OS cells overexpressing CDT1 and CDC6 and depleted for 53BP1. Mean ± SD is indicated by red lines. siLuc CRTL vs siLuc CDT1+CDC6 OE, \*\*\*\**P*<0.0001; siLuc CDT1+CDC6 OE vs siRAD52 CDT1+CDC6 OE, \*\*\*\**P*<0.0001. Statistical analysis was performed using two-tailed Mann-Whitney *U-*tests.

**Extended Data Fig. 8.** High mRNA levels of Cdt1 and Cdc6 are positively correlated with increased copy number of MYC and CCND1 loci in **a,** lung adenocarcinoma **b,** lung squamous cell adenocarcinoma **c,** and head and neck squamous cell carcinoma.

