## Supplementary Material for "Aberrant replication licensing drives Copy Number Gains across species"

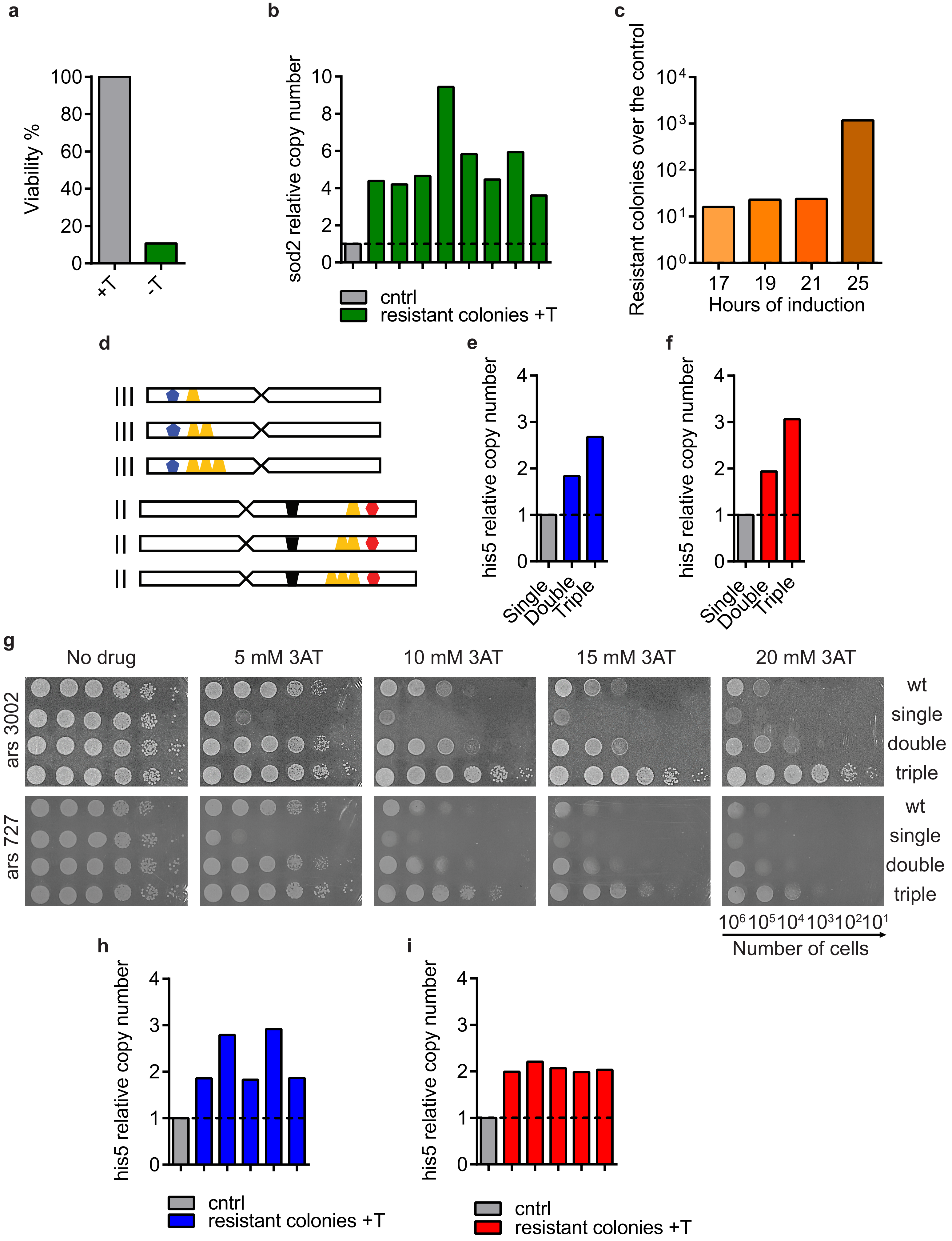
a
Viability %
100
80
60
40
20
0
+T
-T
b
sod2 relative copy number
10
8
6
4
2
0
cntrl
resistant colonies +T
c
Resistant colonies over the control
$10^4$
$10^3$
$10^2$
$10^1$
$10^0$
17
19
21
25
Hours of induction
d
III
III
III
II
II
II
e
his5 relative copy number
4
3
2
1
0
Single
Double
Triple
f
his5 relative copy number
4
3
2
1
0
Single
Double
Triple
g
No drug
5 mM 3AT
10 mM 3AT
15 mM 3AT
20 mM 3AT
ars 3002
ars 727
wt
single
double
triple
wt
single
double
triple
$10^6$ $10^5$ $10^4$ $10^3$ $10^2$ $10^1$
Number of cells
h
his5 relative copy number
4
3
2
1
0
cntrl
resistant colonies +T
i
his5 relative copy number
4
3
2
1
0
cntrl
resistant colonies +T

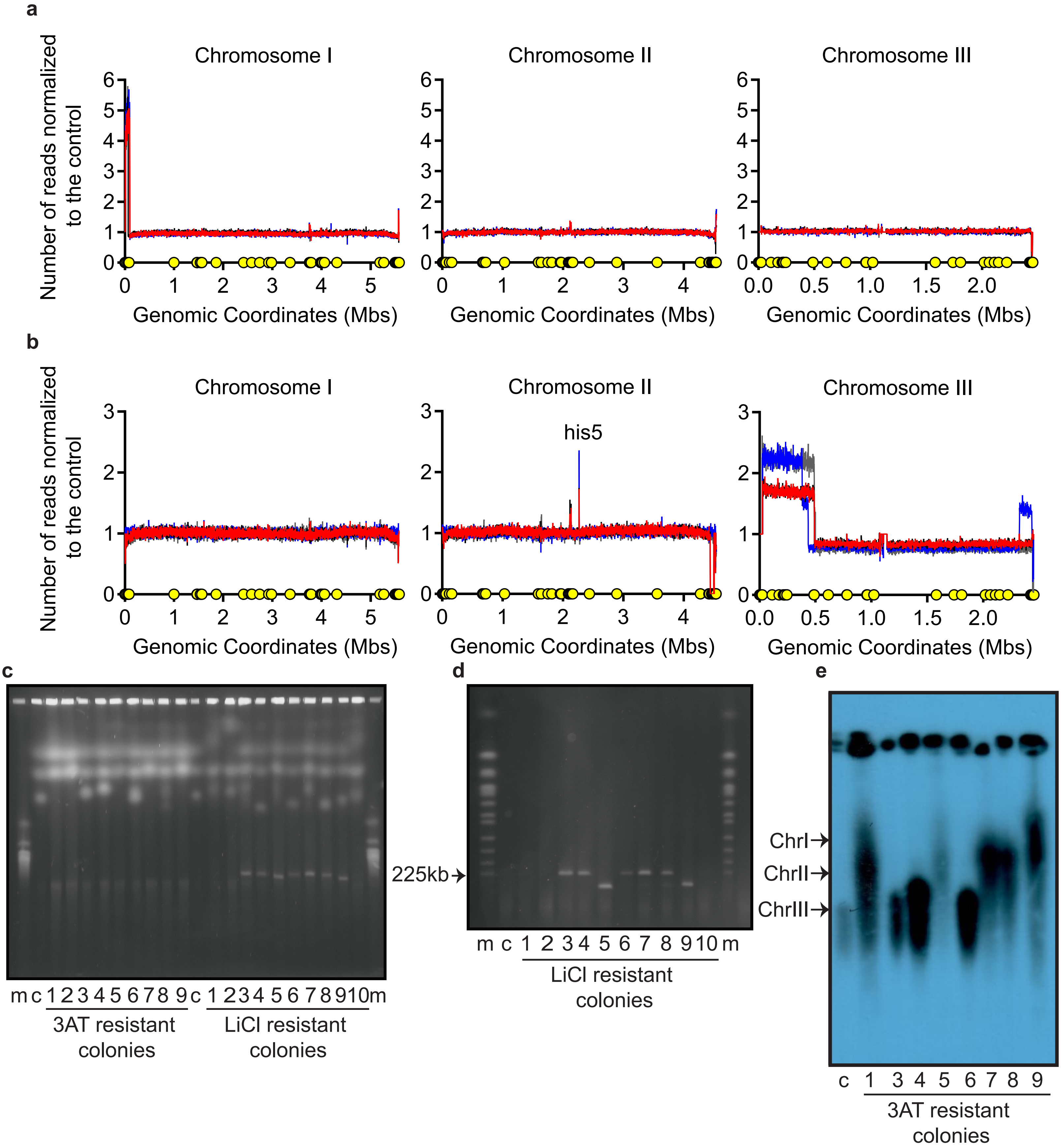
a
Chromosome I
Chromosome II
Chromosome III
Number of reads normalized to the control
Genomic Coordinates (Mbs)
b
Chromosome I
Chromosome II
his5
Chromosome III
Number of reads normalized to the control
Genomic Coordinates (Mbs)
c
m c 1 2 3 4 5 6 7 8 9 c 1 2 3 4 5 6 7 8 9 10 m
3AT resistant colonies
LiCl resistant colonies
225kb
d
m c 1 2 3 4 5 6 7 8 9 10 m
LiCl resistant colonies
e
ChrI
ChrII
ChrIII
c 1 3 4 5 6 7 8 9
3AT resistant colonies

Extended Data Figure 3

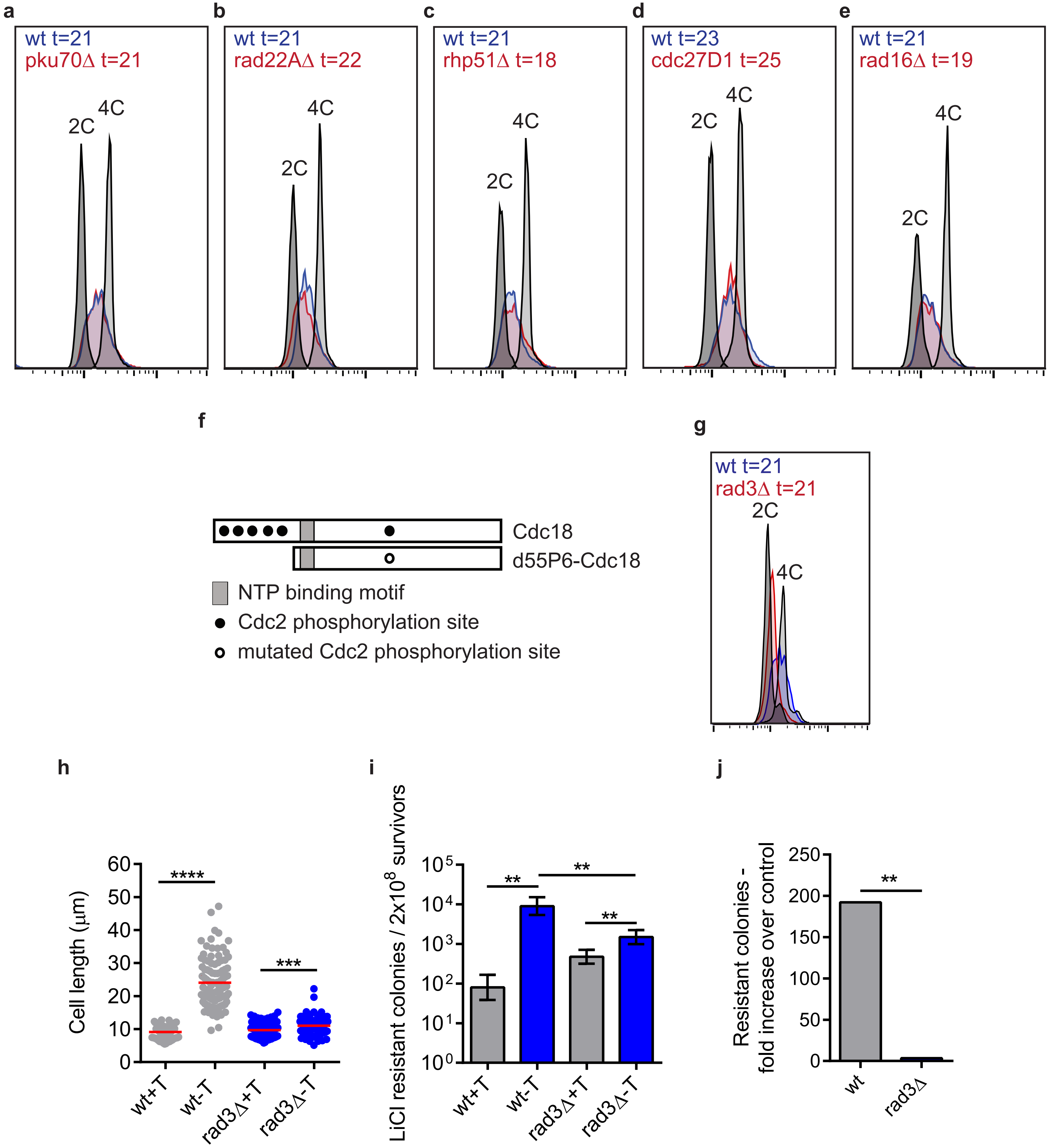

Extended Data Figure 4

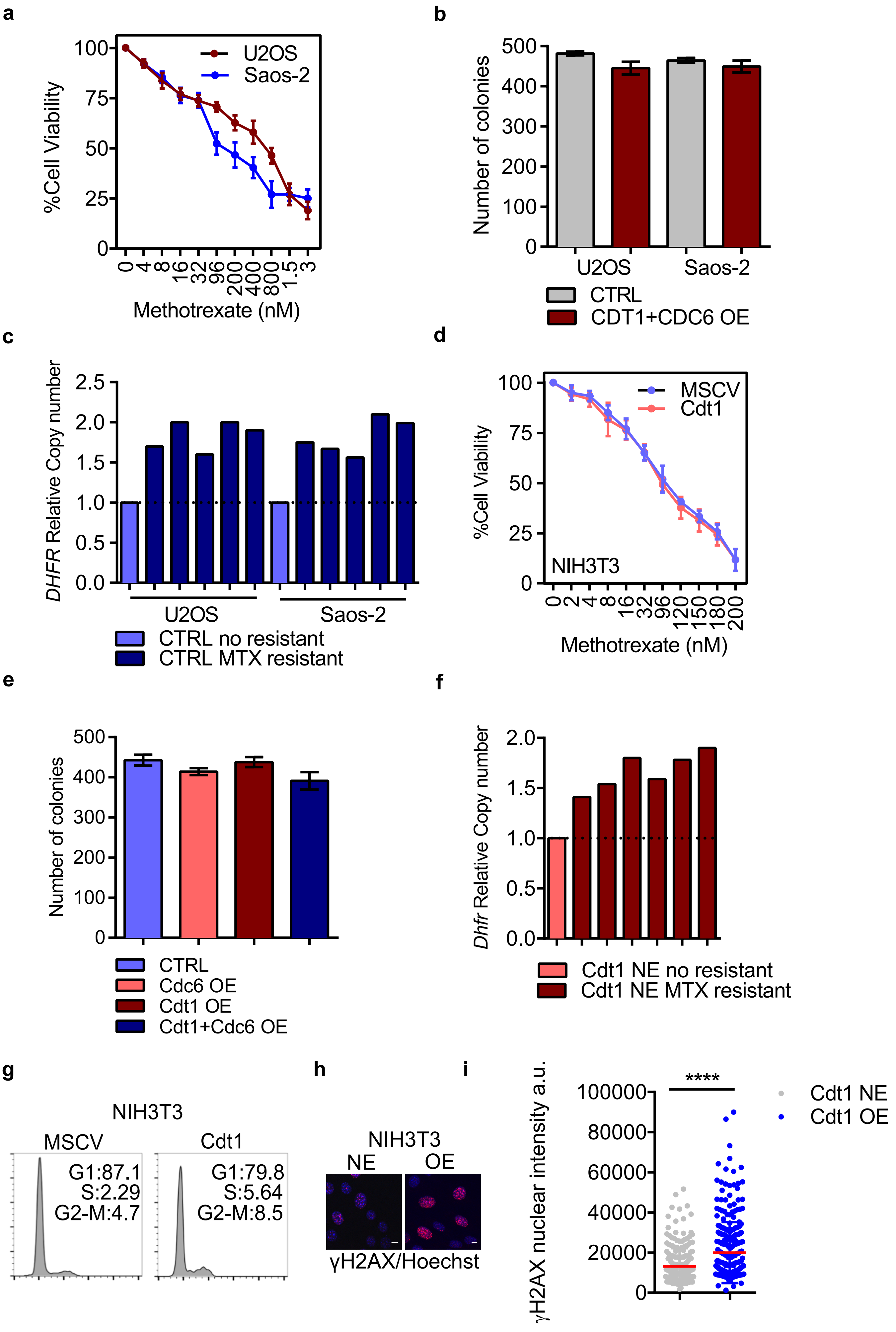

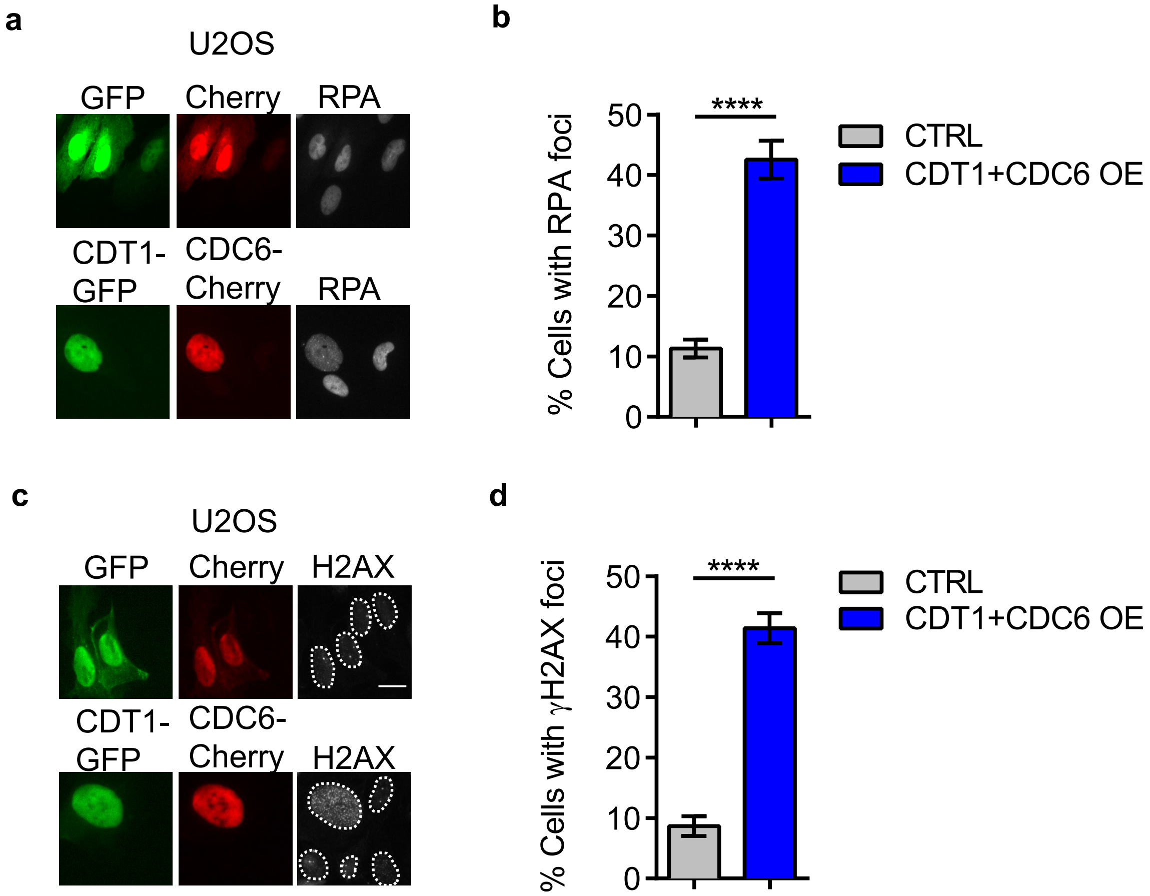
a
U2OS
GFP
Cherry
RPA
CDT1-GFP
CDC6-Cherry
RPA
b
% Cells with RPA foci
****
CTRL
CDT1+CDC6 OE
c
U2OS
GFP
Cherry
H2AX
CDT1-GFP
CDC6-Cherry
H2AX
d
% Cells with γH2AX foci
****
CTRL
CDT1+CDC6 OE

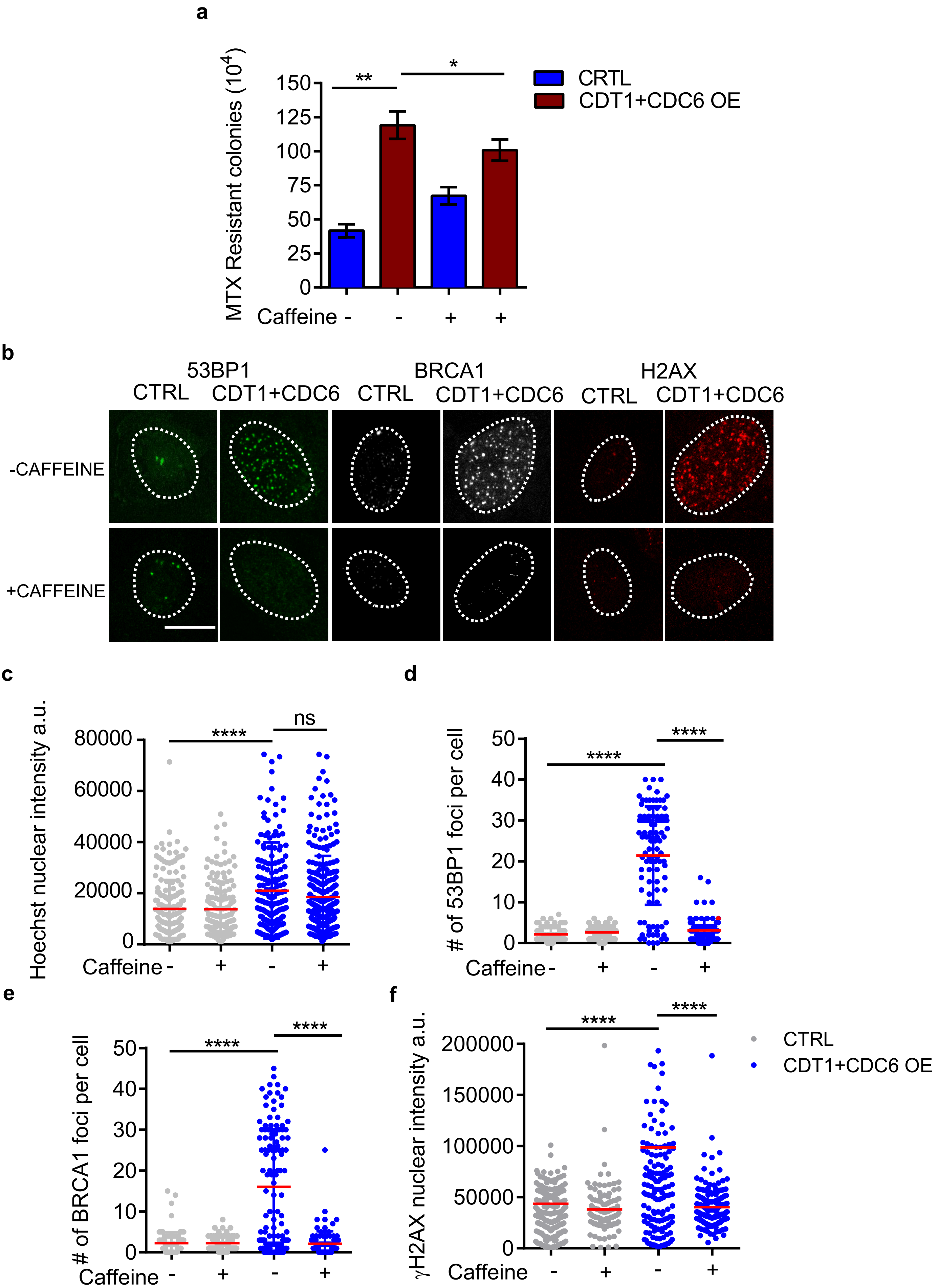
a
MTX Resistant colonies ($10^4$)
150
125
100
75
50
25
0
**
*
CRTL
CDT1+CDC6 OE
Caffeine - - + +
b
53BP1
BRCA1
H2AX
CTRL
CDT1+CDC6
-CAFFEINE
+CAFFEINE
c
Hoechst nuclear intensity a.u.
80000
60000
40000
20000
0
****
ns
Caffeine - + - +
d
### of 53BP1 foci per cell
50
40
30
20
10
0
****
****
Caffeine - + - +
e
### of BRCA1 foci per cell
50
40
30
20
10
0
****
****
Caffeine - + - +
f
γH2AX nuclear intensity a.u.
200000
150000
100000
50000
0
****
****
Caffeine - + - +
CTRL
CDT1+CDC6 OE

Extended Data Figure 7

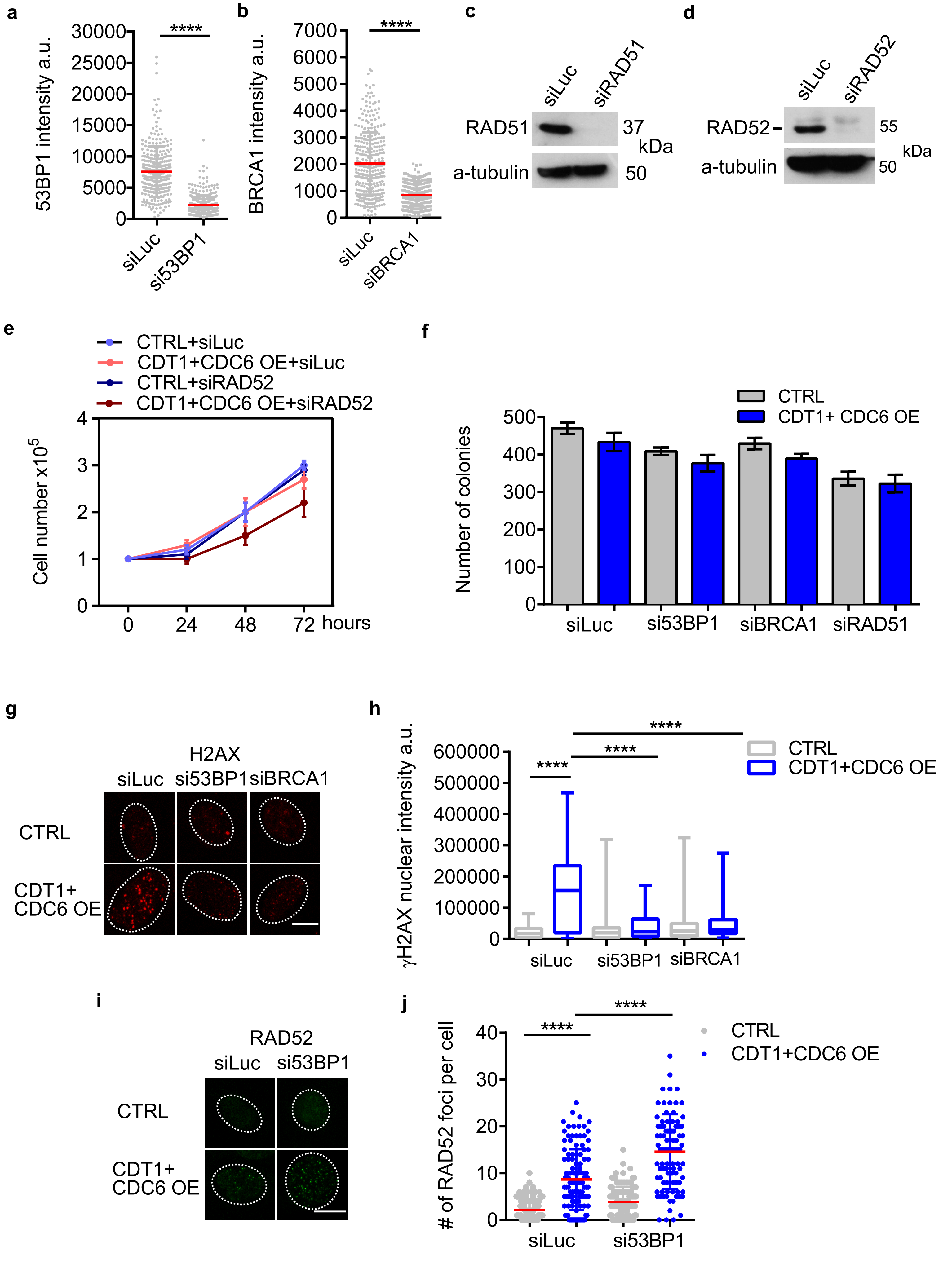

**Extended Data Fig.7c**

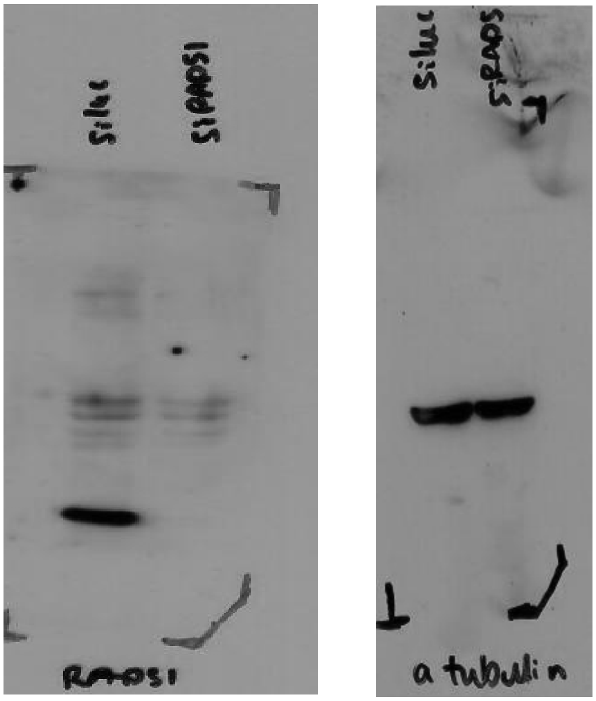


**Extended Data Fig.7d**

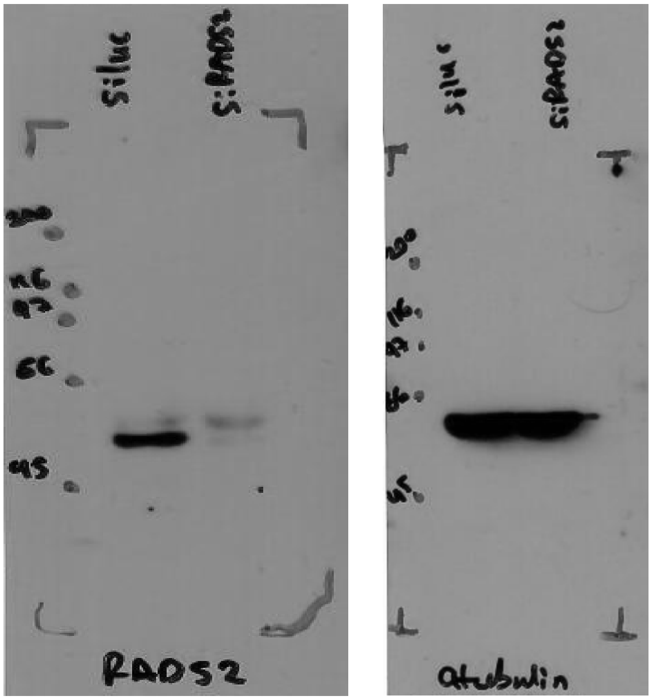

Extended Data Figure 8

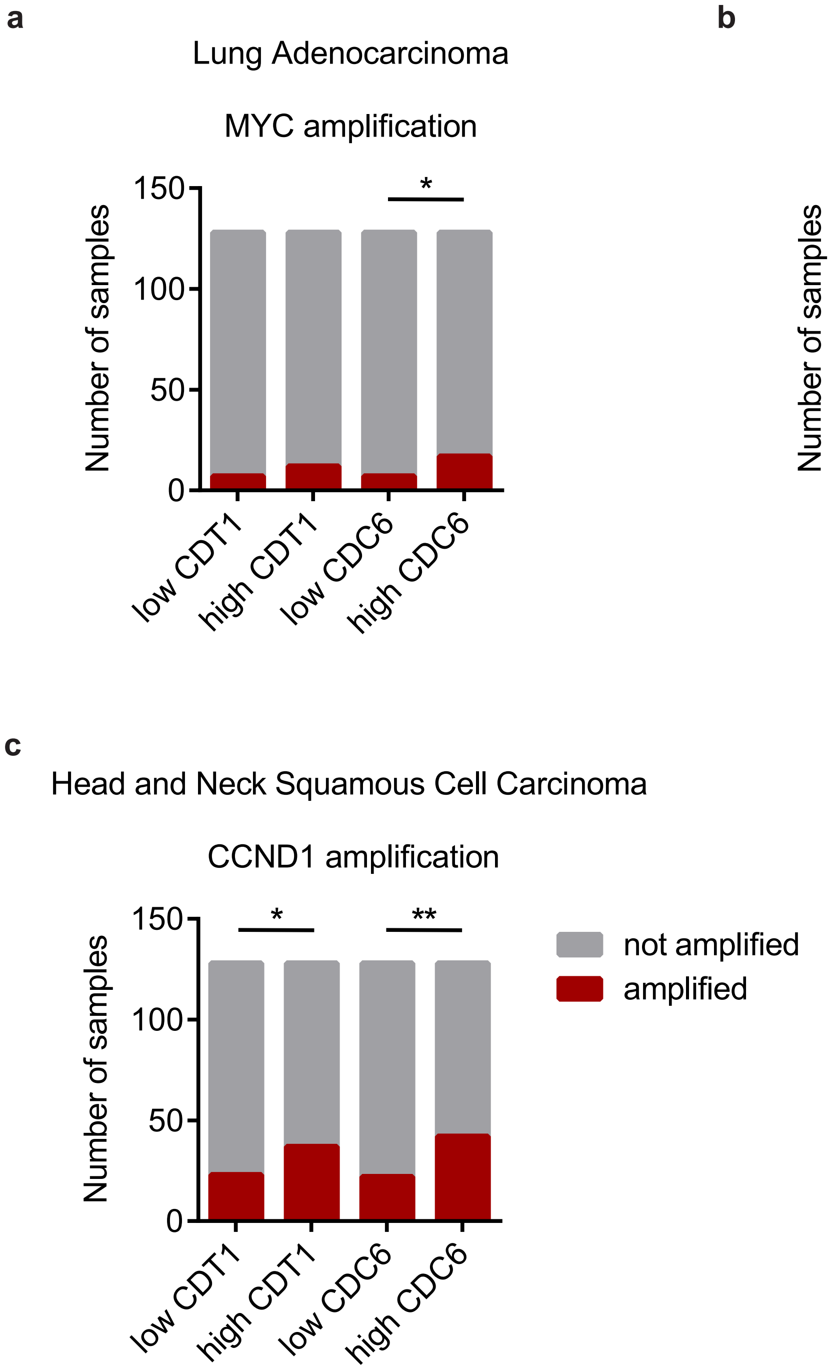

**Supplementary Table 1.** The genotypes and the source of the strains used in this study are presented below.

| Table S1 | | |
|---|---|---|
| **Name** | **Genotype** | **Source** |
| ZP6 | h+, his3-D1, leu1-32, ura4D18, adeM216 | ZL lab |
| ZP7 | h-, his3-D1, leu1-32, ura4D18, adeM210 | ZL lab |
| ZP 151 | h-, nmt3x-d55P6:leu1+, leu1-32, ura4D18 | EMBO J. 1998, 17:5689–98 |
| ZP 153 | h-, nmt1-cdc18:leu1+, leu1-32, uar4D18 | EMBO J. 1998, 17:5689–98 |
| ZP376 | h-, nmt3x-d55P6-leu1+, his5::ura4+ leu1-32, ura4D18 | this study |
| ZP 408 | h-, nmt3x-d55P6-leu1+, his5::ura4+, leu1-32, ura4D18, his5:kan:ars3002 | this study |
| ZP 409 | h-, nmt3x-d55P6-leu1+, his5::ura4+, leu1-32, ura4D18, his5x2:kan:ars3002 | this study |
| ZP 410 | h-, nmt3x-d55P6-leu1+, his5::ura4+, leu1-32, ura4D18, his5x3:kan:ars3002 | this study |
| ZP 466 | h-, nmt3x-d55P6-leu1+, his5::ura4+, leu1-32, ura4D18, his5:kan:ars727 | this study |
| ZP 456 | h-, nmt3x-d55P6-leu1+, his5::ura4+, leu1-32, ura4D18, his5x2:kan:ars727 | this study |
| ZP 467 | h-, nmt3x-d55P6-leu1+, his5::ura4+, leu1-32, ura4D18, his5x3:kan:ars727 | this study |
| ZP 490 | h+, leu1-32, ura4D18, ade6M210, pxd1::kan | Nature Biotechnology 2006, 24:841-845 |
| ZP 491 | h+, leu1-32, ura4D18, ade6M210, rad16::kan | Nature Biotechnology 2006, 24:841-846 |
| ZP 492 | h+, leu1-32, ura4D18, ade6M210, pku70::kan | Nature Biotechnology 2006, 24:841-847 |
| ZP 515 | h+, leu1-32, ura4D18, ade6M210, rad22A::kan | Nature Biotechnology 2006, 24:841-848 |
| ZP 518 | h+, leu1-32, ura4D18, ade6M210, rhp51::kan | Nature Biotechnology 2006, 24:841-849 |
| ZP 450 | h+, leu1-32, ura4D18, ade6M210, rad3::kan | Nature Biotechnology 2006, 24:841-849 |
| ZP 524 | nmt3x-cdc18 d55P6:leu1+, his5::ura4+, leu1-32, ura4D18, ade6M210, his5:kan:ars3002, pxd1::kan | this study |
| ZP 558 | nmt3x-cdc18 d55P6:leu1+, his5::ura4+, leu1-32, ura4D18, his5:ars3002, rad16::kan | this study |
| ZP 535 | nmt3x-cdc18 d55P6:leu1+, his5::ura4+, leu1-32, ura4D18, his5:kan:ars3002, pku70::kan | this study |
| ZP 545 | nmt3x-cdc18 d55P6:leu1+, his5::ura4+, leu1-32, ura4D18, ade6M210, his5:kan:ars3002,rad22A::kan | this study |
| ZP 553 | nmt3x-d55P6:leu1+, his5::ura4+, leu1-32, ura4D18, his5:ars3002, rhp51::kan | this study |
| ZP 481 | nmt3x-cdc18 d55P6-leu1+, leu1-32, ura4D18, rad3::kan | this study |
| ZP 500 | nmt1-cdc18:leu1+, leu1-32, ura4D18, rad3::kan | this study |

**Supplementary Table 2.** The exact sequence of the oligos and the application used are presented below:

| Table S2 | | | |
|---|---|---|---|
| **Number** | **Sequence** | **Application** | |
| 553 | 5’ATTACTTTTTATATGCCTAATAAAAAA GCCATAGTTTAATCTATAGATAACTTTT TTTCCAGTGCACTAACGGACGTTACAC TCATACTCTTCCTTTTTCAATAGC 3’ | his5 disruption using pura4script | use with 554 |
| 554 | 5’GTATTCAATCGTTCCATGGTATAAACA ATAAATACACCAAGGATAAAAAATAAGA AATCAGCGTTACTTAAAAGTATTTACAAA AATTTAACGCGAATTTTAA 3’ | his5 disruption using pura4script | |
| 575 | 5' TCAGGCATCTACGTACAAAGTAAA 3' | check his5 disruption (fw flanking his5) | use with 576 |
| 576 | 5' TGACATTGAATAAGAAAAGAGTGAA 3' | check his5 disruption (ura4 rv) | |
| 577 | 5' TTTTTCTCCATTAAGTAACAAATTCC 3' | check his5 disruption (rv flanking his5) | use with 578 |
| 578 | 5' CGAGAAGCAGAGAGCAGTCA 3' | check his5 disruption (ura4 fw) | |
| 628 | 5’ GGTGATGACGGTGAAAACCT 3’ | check integration in ars3002 | use with 631 |
| 631 | 5’ CTGCTTTGATACATGCACGA 3’ | check integration in ars3002 | |
| 626 | 5’ TGGCGCTAAACAATCTCTGA 3’ | check integration in ars3002 | use with 630 |
| 630 | 5’ AAAGCAAGTCAACACCCCATA 3’ | check integration in ars3002 | |
| 679 | 5’ AAAGTCACGGGTATTAGTGTAACAGA 3’ | check integration in ars727 | use with 680 |
| 680 | 5’ GGTAAAGCATTTGGGGGTCT 3’ | check integration in ars727 | |
| 681 | 5’ CGCCAGCTGAAGCTTCGTA 3’ | check integration in ars727 | use with 682 |
| 682 | 5’ TGTATGGGAATACCGGTGGA 3’ | check integration in ars727 | |
| 695 | 5’ TTCGGAGGCTAGACAAATGG 3’ | real time PCR for ars727 (control region) | use with 696 |
| 696 | 5’ GGGAAATGATCGGGAAAACT 3’ | real time PCR for ars727 (control region) | |
| 701 | 5’ ATGACCATCATCGTGCTGAA 3’ | real time PCR for his5 locus | use with 702 |
| 702 | 5’ ACAACACTCCCTTCGTGCTT 3’ | real time PCR for his5 locus | |
| 727 | 5’ TTGATCGCTGAGTCTGGATG 3’ | real time PCR for sod2 locus | use with 728 |
| 728 | 5’ GCATTCCTCGATGGCTTAAC 3’ | real time PCR for sod2 locus | |
| 795 | 5’ CGTCTGTGAGGGGAGCGTTT 3’ | check deletion of DDR mutants (anneals in the kanamycin cassette) | |
| 796 | 5’ AGAATTGGGTTATTTTCATGTTTGG 3’ | check deletion of rad22A | use with 795 |
| 798 | 5’ AATAGATCACAAGGAAAACTCACCG 3’ | check deletion of pku70 | use with 795 |
| 799 | 5’ GATCCGCCTAAATTCCTAAGAAGGA 3’ | check deletion of rad16 | use with 795 |
| 800 | 5’ GAAGAATTGGGTGTTAAAGCGCATA 3’ | check deletion of pxd1 | use with 795 |
| 828 | 5’ATAACGTACGGTGATTTCAATACCC 3’ | check deletion of rhp51 | use with 795 |
| 852 | 5’ TGCAACCTCGTGTCTTCTTG 3’ | amplicon probe for Southern | use with 853 |
| 853 | 5’ AATGCTTACGGCGATAAACG 3’ | amplicon probe for Southern | |

| | | | |
|---|---|---|---|
| 854 | 5' GGACAATTCCCCAACCTTTT 3' | rDNA probe for Southern | use with 855 |
| 855 | 5' GCAAACCAAAAGGGAGAACA 3' | rDNA probe for Southern | |
| 856 | 5' CAGAGCAATCGCAATGAAAA 3' | telomeric probe for Southern | use with 857 |
| 857 | 5' CGTTCCCGTTAGGAAATCAA 3' | telomeric probe for Southern | |
| 364 | 5′ GGTGAGATTTTGGCGGAGAC 3′ | real time PCR for hDHFR | use with 365 |
| 365 | 5′ TCCCAAGAAAGTAGTGCCCT 3′ | real time PCR for hDHFR | |
| 368 | 5′ CCCAACTTTCCCGCCTCTC 3′ | real time PCR hGAPDH | use with 369 |
| 369 | 5′ CAGCCGCCTGGTTCAA 3′ | real time PCR hGAPDH | |
| 703 | 5′ CAGGCCACCTCAGACTCTTT 3′ | real time PCR for mDHFR | use with 704 |
| 704 | 5′ CAGGCCACCTCAGACTCTTT 3′ | real time PCR for mDHFR | |
| 532 | 5′ TCCCTGGTTAAGCAGTACAG 3′ | real time PCR mHPRT | use with 533 |
| 533 | 5′ GCTTTGTATTTGGCTTTTCC 3′ | real time PCR mHPRT | |
